# Dispensability of the second Cys for phycoviolobilin formation by unusual ring D fixation in the cyanobacteriochrome

**DOI:** 10.1101/2020.12.27.424465

**Authors:** Keiji Fushimi, Rei Narikawa

**Affiliations:** Graduate School of Integrated Science and Technology, Shizuoka University, 836 Ohya, Suruga, Shizuoka 422-8529, Japan; Core Research for Evolutional Science and Technology, Japan Science and Technology; Research Institute of Green Science and Technology, Shizuoka University, 836 Ohya, Suruga, Shizuoka 422-8529, Japan

## Abstract

Cyanobacteriochromes are linear tetrapyrrole-binding photoreceptors produced by cyanobacteria. Their chromophore-binding GAF domains are categorized into many lineages. Among them, the DXCF cyanobacteriochrome GAF domains have a “second Cys” within the DXCF motif in addition to a highly conserved “first Cys” stably ligated to C3^1^ of the A-ring. It has been long known that the second Cys is crucial for two color-tuning events: isomerization activity (reduction of C4=C5 double bond) from the initially incorporated phycocyanobilin to phycoviolobilin and reversible ligation activity to the C10 of the chromophore. Comprehensive site-directed mutagenesis, however, revealed that the second Cys is dispensable for isomerization activity, in which three residues participate by fixing the C- and D-rings. Fixation of the chromophore on both sides of the C5 bridge is necessary, even though one side of the fixation site is far from this bridge, with the other side at C3^1^ fixed by the first Cys.

## Introduction

Cyanobacteriochromes (CBCRs) are linear tetrapyrrole (bilin) -binding photoreceptors that, to date, have been found only in cyanobacteria (Fushimi and Narikawa 2019). The CBCRs are distantly related to red/far-red reversible phytochromes. Only the GAF domain of CBCRs is needed for chromophore incorporation. Phycocyanobilin (PCB) and phycoviolobilin (PVB) are the major chromophores of the CBCR GAF domains (Fig. S1A, B)(Narikawa et al. 2008; Hirose et al. 2008; Ishizuka et al. 2007), whereas biliverdin and 18^1^,18^2^-dihidrobiliverdin can serve as the functional chromophores for some CBCR GAF domains (Moreno et al. 2020; Narikawa et al. 2015; Miyake et al. 2020). A conserved Cys, called the “first Cys” or “canonical Cys,” in the CBCR GAF domain forms a covalent bond with the C3 side chain of the linear tetrapyrrole chromophore via a thioether linkage. The CBCR GAF domains’ light-sensing characteristics depend on the protein and chromophore structures, and absorption of a specific wavelength of light induces *Z/E* isomerization at the C15=C16 double bond to trigger reversible photoconversion between a 15*Z* dark state and a 15*E* photoproduct state (Fig. S1A, B)(Fushimi and Narikawa 2019). The CBCR GAF domains are highly diversified and categorized into many lineages, and consequently, they have various spectral properties.

Of the diverse CBCR GAF domains derived from multiple lineages, DXCF CBCR GAF domains have another conserved Cys, called the “second Cys,” within the Asp-Xaa-Cys-Phe (DXCF) motif (Yoshihara et al. 2004; Ishizuka et al. 2006; Rockwell et al. 2008; Ma et al. 2012; Rockwell, Martin, and Lagarias 2012; Rockwell et al. 2012; Cho et al. 2015; Hasegawa et al. 2018; Narikawa et al. 2011; Song et al. 2011; Narikawa, Kohchi, and Ikeuchi 2008; Enomoto et al. 2012). Typical DXCF CBCR GAF domains initially incorporate PCB as a precursor chromophore, which is then isomerized to PVB (PCB-to-PVB isomerization activity)(Ishizuka et al. 2007, 2011). The second Cys contributes to this isomerization activity. In addition, the second Cys reversibly attaches to or detaches from the C10 position of PVB during the photoconversion process (i.e., reversible ligation activity)(Rockwell et al. 2008; Ishizuka et al. 2011; Burgie et al. 2013; Cornilescu et al. 2014; Narikawa et al. 2013). In this context, the second Cys residue possesses dual functions involved in isomerization activity and reversible ligation activity.

In our previous study, we firstly identified a novel CBCR GAF domain, AM1_1499g1, which lacks the second Cys, although the domain belongs to a specific DXCF CBCR lineage (AM1_1499g1/AM1_6305g1 lineage)(Fushimi et al. 2020). AM1_1499g1 contains PCB and exhibits an orange/green photocycle but lacks isomerization activity and reversible ligation activity (Fig. S1C). Furthermore, we have succeeded in evolution-inspired color-tuning based on the AM1_1499g1 scaffold; the introduction of the second Cys (S_118_C variant) results in the acquisition of isomerization activity but not reversible ligation activity, with the exhibition of a yellow/teal photocycle (Fig. S1C). In addition, replacement of Tyr and Thr near the D-ring of the chromophore with Leu and Asn, respectively, (S_118_C/Y_151_L/T_159_N variant, based on the S_118_C variant molecule) yields a dark state blue-shift, likely due to the D-ring twist (Fig. S1D, E), with the molecule subsequently displaying a green/teal photocycle (Fig. S1C). Furthermore, replacement of the His next to the first Cys residue with Tyr (S_118_C/H_147_Y variant) enables reversible ligation activity, in which the bulky side chain may push the C10 bridge between the B-ring and C-ring toward the second Cys to facilitate reversible ligation (Fig. S1D, E) with the presentation of a blue/teal photocycle (Fig. S1C).

On the other hand, AM1_6305g1, a close homolog of AM1_1499g1, retains the second Cys, and displays a green/teal photocycle (Hasegawa et al. 2018). AM1_6305g1 can isomerize PCB to PVB but cannot ligate the second Cys to the C10 position. We have performed reverse engineering on AM1_6305g1, based on the engineering work of AM1_1499g1, and succeeded in developing two variant molecules showing yellow/teal and blue/teal photocycles for modification of the dark-state D-ring twist and reversible Cys ligation (Fushimi et al. 2020).

However, the replacement of the second Cys with Ser unexpectedly did not affect the isomerization activity. Starting from this unanticipated observation, we identified three residues key for the isomerization activity in addition to the second Cys in this study. These findings provide a general concept for the isomerization mechanism.

## Results and discussion

### Second Cys residue is dispensable for PCB-to-PVB isomerization in the AM1_6305g1 scaffold

We previously reported that one of the DXCF lineages, AM1_1499g1/AM1_6305g1, exhibits three distinctive photocycles: orange/green, green/teal, and blue/teal. The second Cys of the blue/teal molecules retains the dual function of PCB-to-PVB isomerization activity and reversible ligation activity. In contrast, the second Cys of the green/teal molecules, including AM1_6305g1, lost the reversible ligation activity, and the molecule only displays isomerization activity. The orange/green molecule, AM1_1499g1, has lost the second Cys and does not show isomerization activity nor reversible ligation activity (Fig. S1C, E). Based on this natural diversity, we have engineered AM1_1499g1 to display various photocycles (Fig. S1C)(Fushimi et al. 2020). The introduction of the second Cys in AM1_1499g1, as part of the primary engineering phase, resulted in the acquisition of isomerization activity but not reversible ligation activity (Fig. S1C). This result encouraged us to perform reverse engineering on AM1_6305g1 for loss of the isomerization activity.

We thus replaced the second Cys of AM1_6305g1 with Ser, the original residue in AM1_1499g1 (AM1_6305g1_C_99_S) (Fig. S2A-i). Unexpectedly, AM1_6305g1_C_99_S showed a green/teal photocycle, which is nearly the same as that of the wild-type (Fig. 1A, Fig. 2A, and Table 1). Furthermore, the chromophore incorporated into AM1_6305g1_C_99_S was assigned not to PCB but to PVB by comparing the denatured difference absorption spectrum (dark state–photoproduct state) of the C_99_S variant with those of the PCB- and PVB-binding CBCR GAF domains (Fig. 3A and Table 1). These results indicate that the second Cys residue is dispensable for the isomerization activity in the AM1_6305g1 scaffold. Ser and Cys have a common skeleton with distinctive functional groups at the end of the side-chains; a hydroxy group for Ser and a thiol group for Cys. We speculate that the hydroxy group of Ser can complement the functionality of the thiol group of Cys in AM1_6305g1.

**Figure 1.**
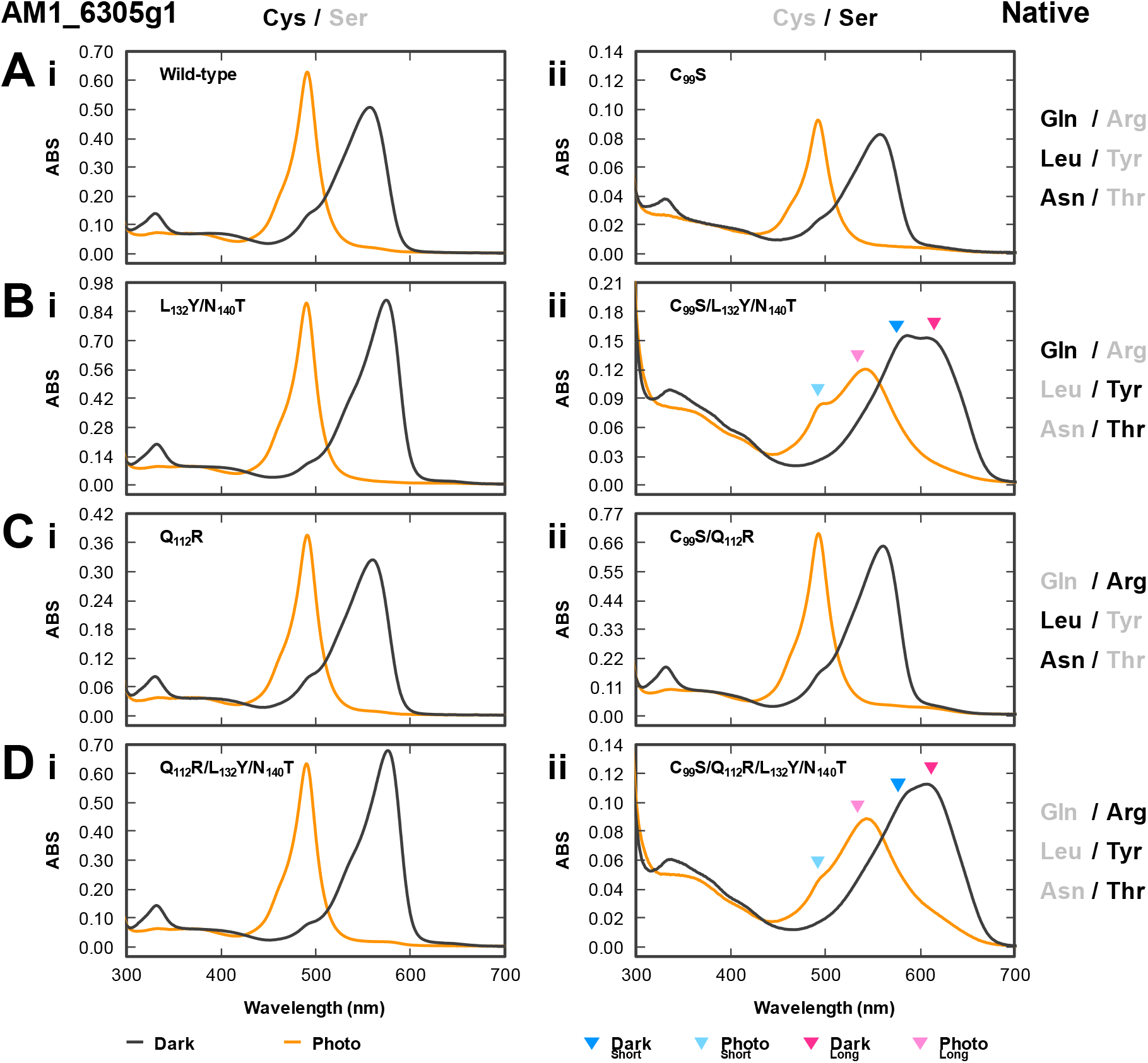
Absorption spectra of native wild-type and variant molecules of the AM1_6305g1 scaffold. (A–D) Absorption spectra of the dark state (solid gray line) and the photoproduct state (solid orange line) were measured at room temperature. In the case of molecules containing two photoconvertible components, both components were fixed to the dark state or the photoproduct state. Wavelength area corresponding to the positive (dark state, deep color triangle) and negative (photoproduct state, light color triangle) peaks of the normalized difference spectra from the two photoconvertible components (yellow/teal component in short wavelength region, cyan; orange/green component in the long-wavelength region, magenta) were assigned by colored triangles. The normalized difference spectra are shown in Figure 2. (A) The molecules having Gln, Leu, and Asn in the Gln/Arg, Leu/Tyr, and Asn/Thr positions, respectively. (B) The molecules having Gln, Tyr, and Thr in these positions. (C) The molecules having Arg, Leu, and Asn in these positions. (D) The molecules having Arg, Tyr, and Thr in these positions. (i, ii) The molecules having Cys (i) or Ser (ii) in the second Cys position.

**Figure 2.**
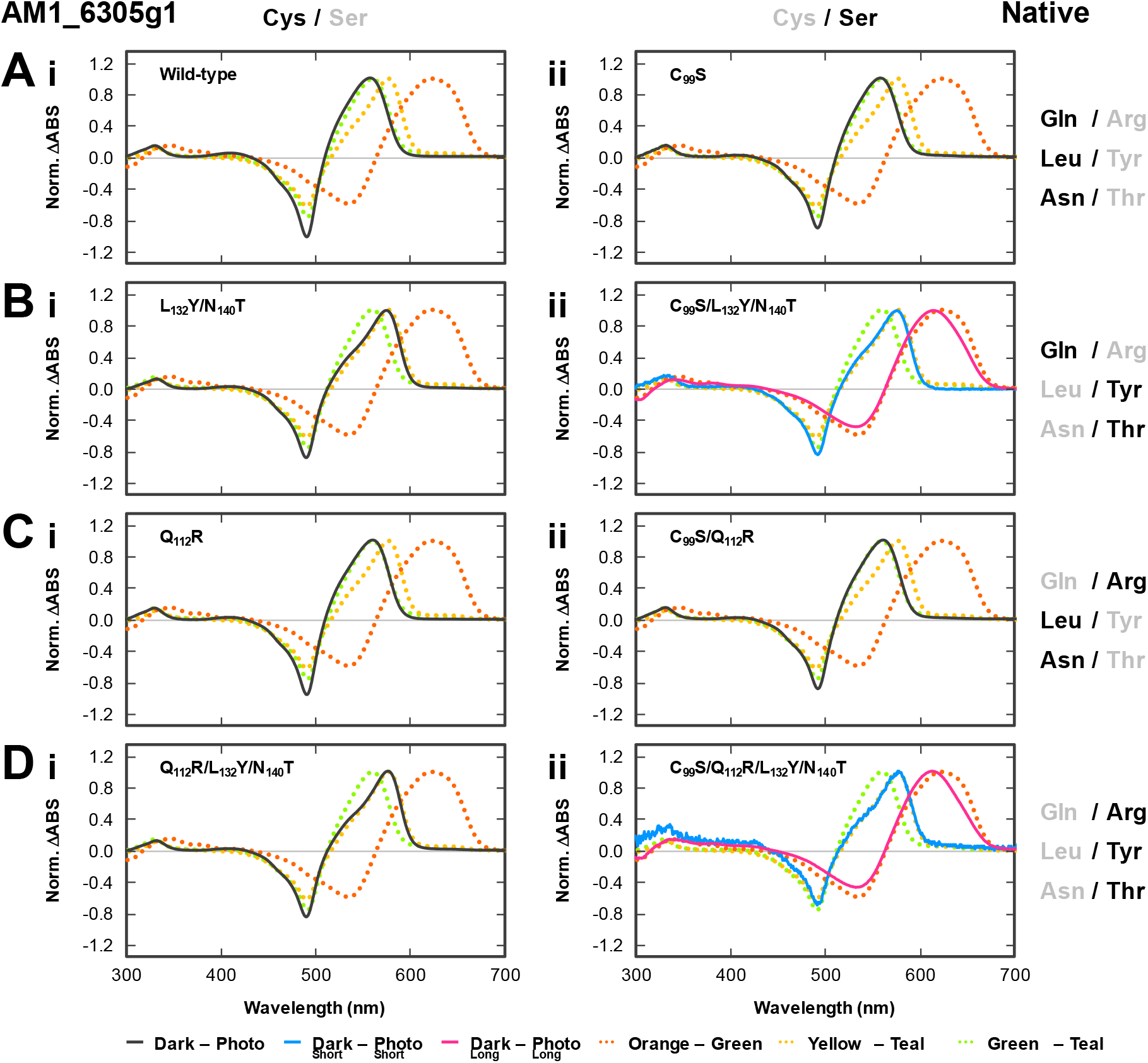
Normalized difference absorption spectra (dark state-photoproduct state) of native wild-type and variant molecules of the AM1_6305g1 scaffold. (A–D) Difference spectra of the molecules containing a single photoconvertible component (solid gray line) were calculated from the native absorption spectra shown in Figure 1. On the other hand, difference spectra of the molecules containing two photoconvertible components (yellow/teal component in the short wavelength region, solid cyan line; orange/green component in the long-wavelength region, solid magenta line) were calculated from the native absorption spectra shown in Figure S5. In Figure S5, we separately excited each photoconvertible component. These spectra were compared with those of the known orange/green, yellow/teal, and green/teal photoconvertible molecules (AM1_1499g1 wild-type, orange dotted line; S_118_C, yellow dotted line; S_118_C/Y_151_L/T_159_N, green dotted line, respectively). These absorption peaks are summarized in Table 1. (A) The molecules having Gln, Leu, and Asn in the Gln/Arg, Leu/Tyr, and Asn/Thr positions, respectively. (B) The molecules having Gln, Tyr, and Thr in these positions. (C) The molecules having Arg, Leu, and Asn in these positions. (D) The molecules having Arg, Tyr, and Thr in these positions. (i, ii) The molecules having Cys (i) or Ser (ii) in the second Cys position.

**Figure 3.**
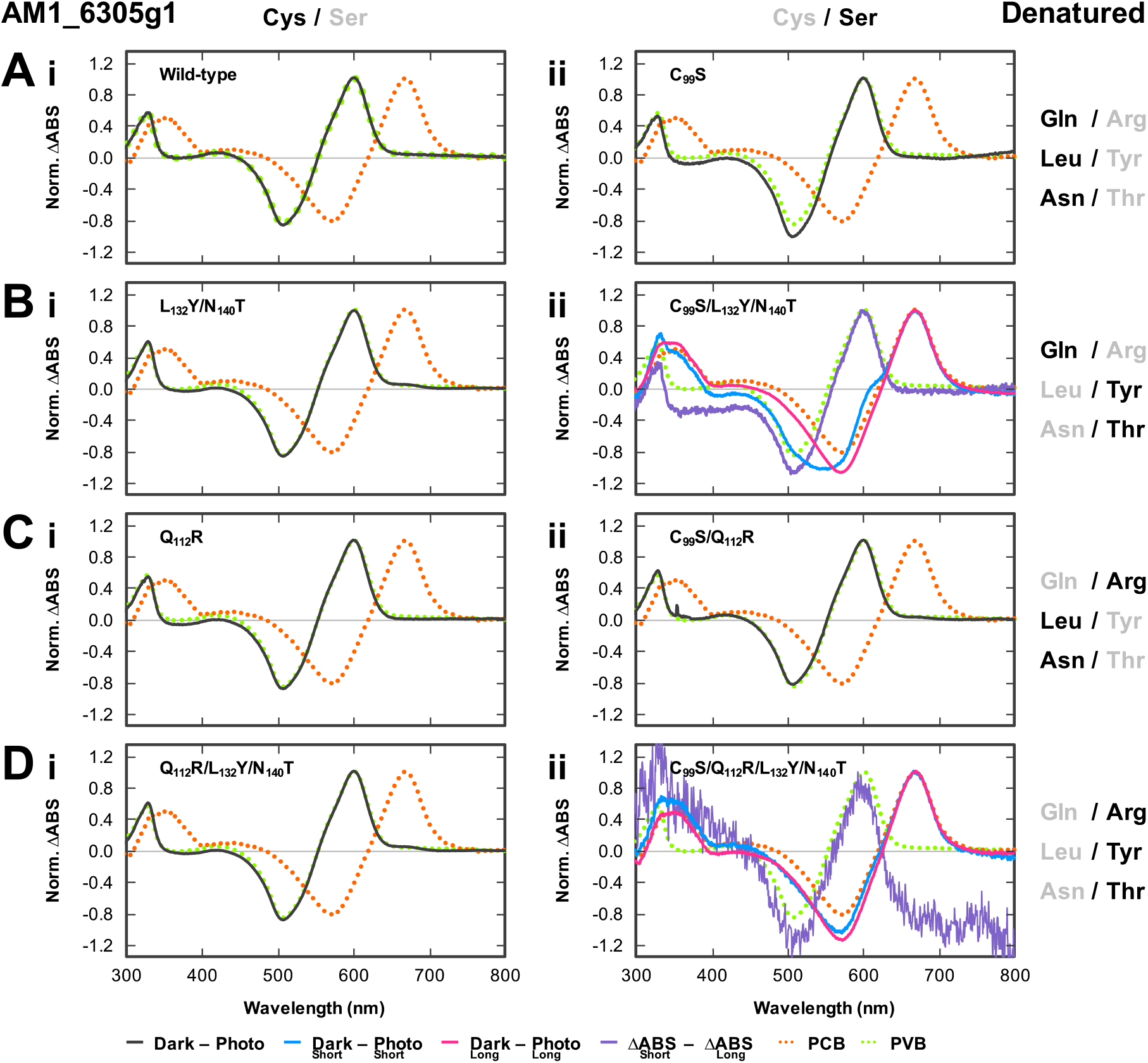
Normalized difference absorption spectra (dark state-photoproduct state) of denatured AM1_6305g1 species. (A–D) Difference spectra of the molecules containing a single photoconvertible component (solid gray line) and two photoconvertible components (yellow/teal component in the short wavelength region, solid cyan line; orange/green component in the long-wavelength region, solid magenta line) were calculated from the denatured absorption spectra shown in Figure S6. Detailed experimental protocol was described in Materials and Methods. Furthermore, the normalized double-difference spectra (yellow/teal photoconvertible component in the short wavelength region, solid violet line) of AM1_6305g1_C_99_S/L_132_Y/N_140_T and C_99_S/Q_112_R/L_132_Y/N_140_T were calculated from these normalized difference absorption spectra. These spectra were compared with those of the known PCB- and PVB-binding molecules (AM1_1499g1 wild-type, orange dotted line; AM1_6305g1 wild-type, green dotted line, respectively). These absorption peaks are summarized in Table 1. (A) The molecules having Gln, Leu, and Asn in the Gln/Arg, Leu/Tyr, and Asn/Thr positions, respectively. (B) The molecules having Gln, Tyr, and Thr in these positions. (C) The molecules having Arg, Leu, and Asn in the positions. (D) The molecules having Arg, Tyr, and Thr in these positions. (i, ii) The molecules having Cys (i) or Ser (ii) in the second Cys position.

**Table 1.**
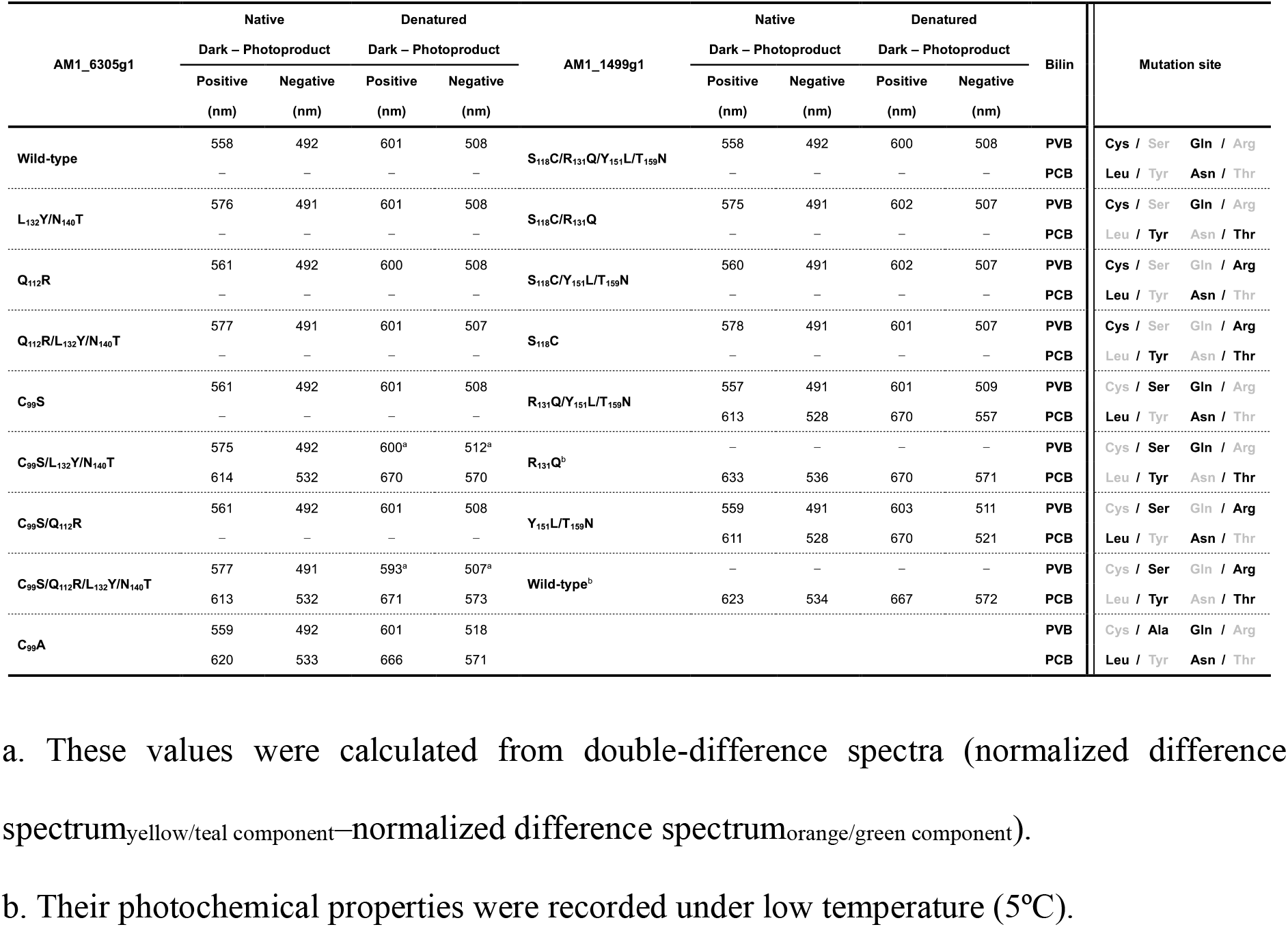
Wavelength peaks of the difference absorption spectra of the AM1_6305g1 and AM1_1499g1 variant molecules under native and denatured conditions (shown in Figures 2, 3, 4, 6, and 7).Figures

We next replaced these polar amino acids with Ala (AM1_6305g1_C_99_A), which has a nonpolar methyl group smaller than that of Ser and Cys to verify this assumption (Fig. S2A-ii). Spectral analysis revealed two photoconvertible components for AM1_6305g1_C_99_A; green/teal and orange/green (Fig. 4A). We were able to separately excite a single component and independently characterize the photoconversion properties and the binding chromophore species based on the absorption spectra of the native and denatured molecules using various monochromic light sources (Fig. 4B, C and Table 1). The green/teal component bound PVB (Fig. 4C and Table 1, shown by a cyan line) and showed reversible photoconversion between a green-absorbing dark state (~560 nm) and a teal-absorbing photoproduct state (~490 nm) (Fig. 4A, shown by cyan arrowheads). The orange/green component bound PCB (Fig. 4C and Table 1, shown by a magenta line) and showed reversible photoconversion between an orange-absorbing dark state (~620 nm) and a green-absorbing photoproduct state (~550 nm) (Fig. 4A, shown by magenta arrowheads). Our findings taken together indicate that AM1_6305g1_C_99_A is a mixture of PCB- and PVB-binding populations.

**Figure 4.**
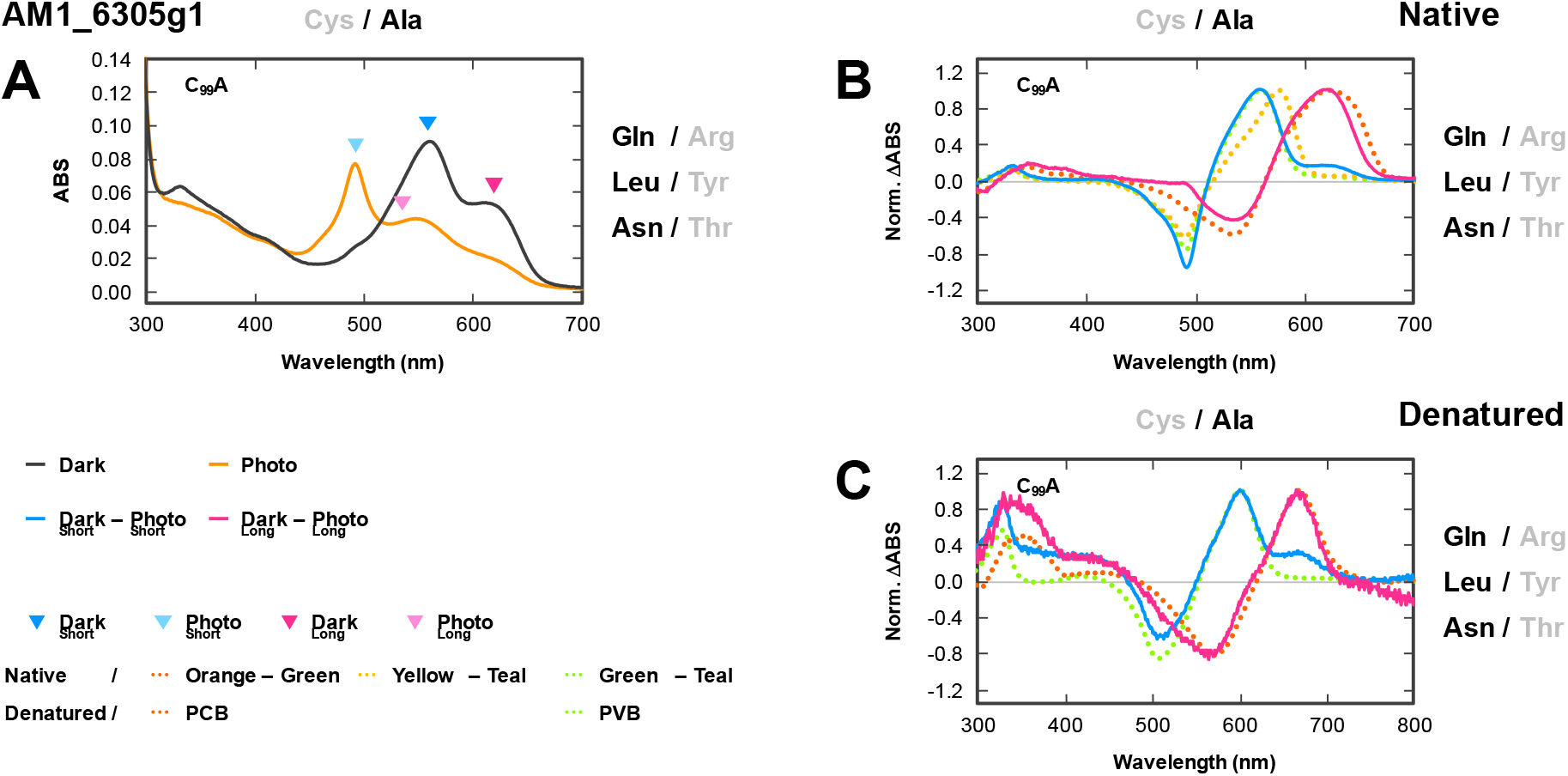
UV-Vis spectra of native and denatured AM1_6305g1_C_99_A. (A) The absorption spectra (dark state, solid gray line; photoproduct state, solid yellow line) of AM1_6305g1_C99A were measured at room temperature. Wavelength area of the positive (dark state, deep color triangle) and negative (photoproduct state, light color triangle) peaks of the normalized difference spectra from the two photoconvertible components (green/teal component in short wavelength region, cyan; orange/green component in the long-wavelength region, magenta) were assigned by colored triangles. (B, C) The normalized difference spectra (dark state–photoproduct state) of the molecules containing the two photoconvertible components (green/teal component in the short wavelength region, cyan solid line; orange/green component in the long-wavelength region, solid magenta line) were calculated from the native (B) and denatured (C) absorption spectra shown in Figure S5 and Figure S6. The native spectra were compared with those of the known orange/green, yellow/teal, and green/teal photoconvertible molecules (AM1_1499g1 wild-type, orange dotted line; S_118_C, yellow dotted line; S_118_C/Y_151_L/T_159_N, green dotted line, respectively). On the other hand, the denatured spectra were compared with those of the known PCB- and PVB-binding molecules (AM1_1499g1 wild-type, orange dotted line; AM1_6305g1 wild-type, green dotted line, respectively). These absorption peaks are summarized in Table 1.

In conclusion, the second Cys residue is dispensable for the isomerization activity in the AM1_6305g1 scaffold. Ser, but not Ala, could complement the function of the Cys residue in the AM1_6305g1 scaffold. Furthermore, the fact that AM1_6305g1_C_99_A showed partial isomerization activity indicates the contribution of another residue(s) to the isomerization activity.

### Leu_132_ and Asn_140_ in AM1_6305g1 are crucial for efficient PCB-to-PVB isomerization without the second Cys residue

We have found that the AM1_6305g1 scaffold is clearly distinct from the AM1_1499g1 scaffold in the context of isomerization activity, despite their close homologous relationship. Namely, the second Cys residue is dispensable for isomerization activity in the AM1_6305g1 scaffold but not in the AM1_1499g1 scaffold, indicating that the distinct residues between these two molecules are determinants for this divergence. We have already determined in a previous study that such amino acid alterations are crucial for color-tuning of the dark state via a D-ring twist; Leu_132_/Asn_140_ in AM1_6305g1 and Tyr_151_/Thr_159_ in AM1_1499g1 (Fig. S1C–E)(Fushimi et al. 2020). We hypothesize that the alterations of these two residues are key not only for the dark-state color-tuning but also for isomerization activity. Although replacement of Leu_132_/Asn_140_ with Tyr/Thr based on the AM1_6305g1 wild-type background (AM1_6305g1_L_132_Y/N_140_T) resulted in a red shift of the dark state, no effects on the isomerization activity have been detected (Fig. 1B-i, Fig. 2B-i, Fig. 3B-i, and Table 1)(Fushimi et al. 2020). In other words, this variant molecule has a single yellow/teal photoconvertible component, and the binding chromophore species of this variant molecule are composed of PVB as well as the wild-type. In this context, these two residues may be crucial for the isomerization activity without the second Cys residue.

To verify this hypothesis, we replaced these two residues (Leu132 and Asn140) with Tyr and Thr, respectively, based on the C_99_S background molecule (AM1_6305g1_C_99_S/L_132_Y/N_140_T) (Fig. S2A-i). AM1_6305g1_C_99_S/L_132_Y/N_140_T had two photoconvertible components displaying yellow/teal and orange/green photocycles, in contrast to the L_132_Y/N_140_T variant (Fig. 1B-ii). The difference spectra of these two components were similar to those of the PVB-binding and PCB-binding reference molecules showing yellow/teal and orange/green photocycles, respectively (Fig. 2B-ii and Table 1). The chromophores incorporated into these photoconvertible components, the yellow/teal and orange/green ones of AM1_6305g1_C_99_S/L_132_Y/N_140_T, were assigned to PVB and PCB, respectively, based on their denatured difference absorption spectra (Fig. 3B-ii and Table 1). We could not obtain a pure difference spectrum of the denatured yellow/teal component because there was much less of the yellow/teal component than the orange/green one. However, a double-difference spectrum (normalized difference spectrum_yellow/teal component_–normalized difference spectrum_orange/green component_) clearly showed that the yellow/teal component was derived from PVB incorporation (Fig. 3B-ii and Table 1, shown by a violet line). These results suggested that the Ser residue itself could not fully complement the function of the second Cys residue and that the Leu and Asn residues, but not the Tyr and Thr residues, could support the Ser residue for full isomerization of PCB to PVB (Fig. 1A, B, Fig. 2A, B, Fig. 3A, B, and Table 1). Thus, these two amino acid positions are involved not only in dark-state color-tuning but also in isomerization activity.

We constructed singly mutated variant molecules based on the C_99_S background molecule (AM1_6305g1_C_99_S/L_132_Y and C_99_S/N_140_T) to elucidate the contribution of each residue to the isomerization activity (Fig. S2A-iii). These two variant molecules had yellow/teal and orange/green components, as well as the doubly mutated ones (AM1_6305g1_C_99_S/L_132_Y/N_140_T) (Fig. S3A, B). A large amount of yellow/teal component was detected for the C_99_S/L_132_Y variant, whereas a smaller amount of the same component was detected for the C_99_S/N_140_T variant; the contribution of the residues at the Leu/Tyr position to the isomerization activity was larger than that of the Asn/Thr position.

The PVB-binding dark state of the AM1_6305g1_C_99_S/L_132_Y/N_140_T, the yellow-absorbing form, was red-shifted in comparison with that of the C_99_S background molecule, which is consistent with the same L_132_Y/N_140_T replacement on the wild-type background (Fig. 1A, B, Fig. 2A, B, and Table 1). This finding could be explained by the previously proposed mechanism, in which replacement with Tyr and Thr resulted in the cancellation of the D-ring twist, leading to the red shift (Fushimi et al. 2020). On the other hand, because the C_99_S background molecule did not contain any PCB-binding component, we could not discuss the dark-state color-tuning of the PCB-binding ones (Fig. 2A-ii and Table 1).

In conclusion, the Ser residue can fully complement the isomerization function of the second Cys residue only when the Leu and Asn residues hold the D-ring with twisted geometry. However, the AM1_6305g1_C_99_S/L_132_Y/N_140_T variant still contained PVB-binding yellow/teal components, albeit at lower levels, indicating the additional contribution of another residue(s).

### Gln_112_ in AM1_6305g1 is a third key residue for isomerization activity in the absence of the second Cys residue

In the previous section, we focused on the Leu/Tyr and Asn/Thr positions in the context of residues specific to AM1_1499g1 among the molecules within the AM1_1499g1/AM1_6305g1 lineage (Fig. S1E). We next focused on the residue(s) specific to AM1_6305g1 near the chromophore among the molecules of this lineage and found the Gln/Arg position near the C-ring propionate. AM1_6305g1 specifically possesses a Gln residue (Gln112) at this position, whereas other molecules within this lineage contain Arg residues (Fig. S1D, E). We constructed four variants (AM1_6305g1_Q_112_R, C_99_S/Q_112_R, Q_112_R/L_132_Y/N_140_T, and C_99_S/Q_112_R/L_132_Y/N_140_T) based on their corresponding background molecules (AM1_6305g1 wild-type, C_99_S, L_132_Y/N_140_T, and C_99_S/L_132_Y/N_140_T), respectively, to elucidate the function of the Gln residue (Fig. S2A-i). A single green/teal component was detected for the Q_112_R and C_99_S/Q_112_R variants as well as their background molecules (wild-type and C_99_S) (Fig. 1C, Fig. 2C, and Table 1), whereas a single yellow/teal component was detected for the Q_112_R/L_132_Y/N_140_T variant as well as its background molecule (L_132_Y/N_140_T) (Fig. 1D-i, Fig. 2D-i, and Table 1). The chromophores incorporated into these variant molecules were assigned to PVB, indicating that they retained full isomerization activity (Fig. 3C, 3D-i, and Table 1). Conversely, the C_99_S/Q_112_R/L_132_Y/N_140_T variant molecule had close to a single orange/green component with a faint yellow/teal component, which corresponded to the PCB- and PVB-binding populations, respectively (Fig. 1D-ii, Fig. 2D-ii, Fig. 3D-ii, and Table 1). In conclusion, the replacement of a total of four residues (C_99_S/Q_112_R/L_132_Y/N_140_T) resulted in almost complete inactivation of the isomerization activity.

### Summary of engineering on AM1_6305g1

Based on the sequence comparison, we have identified four key residues (Cys_99_/Gln_112_/Leu_132_/Asn_140_) of the AM1_6305g1 involved in the isomerization activity from PCB to PVB (Fig. S1D, E). Only the second Cys (Cys_99_) is sufficient for full isomerization among these residues, while any replacement of the other residues (Gln_112_Arg, Leu_132_Tyr and Asn_140_Thr) under the presence of Cys_99_ did not at all affect the isomerization activity (Fig. 1A-i to D-i, Fig. 2A-i to D-i, Fig. 3A-i to D-i, and Table 1). Leu_132_ and Asn_140_ plays a central role in isomerization activity under the presence of a Ser in place of the second Cys, while Gln_112_ plays a supportive role (Fig. 1A-ii to D-ii, Fig. 2A-ii to D-ii, Fig. 3A-ii to D-ii, and Table 1). Namely, AM1_6305g1 has constructed a robust system for isomerization activity, in which Gln_112_, Leu_132_ and Asn_140_ can support the Ser residue, substitute of the second Cys.

### Reverse engineering on AM1_1499g1

In contrast to AM1_6305g1, AM1_1499g1 has Ser_118_/Arg_131_/Tyr_151_/Thr_159_ residues corresponding to Cys_99_/Gln_112_/Leu_132_/Asn_140_ of AM1_6305g1 (Fig. S1D, E). This fact suggests that AM1_1499g1 has completely lost the redundancy system for the isomerization activity observed for AM1_6305g1. To verify this hypothesis, we performed reverse engineering on AM1_1499g1 to generate variant molecules (Y_151_L/T_159_N, R_131_Q, R_131_Q/Y_151_L/T_159_N, S_118_C, S_118_C/Y_151_L/T_159_N, S_118_C/R_131_Q, and S_118_C/R_131_Q/Y_151_L/T_159_N) based on the wild-type molecule corresponding to each AM1_6305g1 variant (Fig. S2B-i) and compared each (Figs. 5–7, and Table 1). All variant molecules possessing the second Cys (S_118_C, S_118_C/Y_151_L/T_159_N, S_118_C/R_131_Q, and S_118_C/R_131_Q/Y_151_L/T_159_N) incorporated only PVB and showed yellow/teal or green/teal reversible photoconversion (Fig. 5A-i to D-i, Fig. 6A-i to D-i, Fig. 7A-i to D-i, and Table 1). This result is fully consistent with the case of AM1_6305g1; the second Cys is also sufficient for the isomerization activity for the AM1_1499g1 scaffold. Variants possessing Tyr_151_/Thr_159_ showed a yellow/teal photocycle, while variants possessing Leu_151_/Asn_159_ showed a green/teal photocycle, which is well explained by the dark-state trapped twist model (Fushimi et al. 2020).

**Figure 5.**
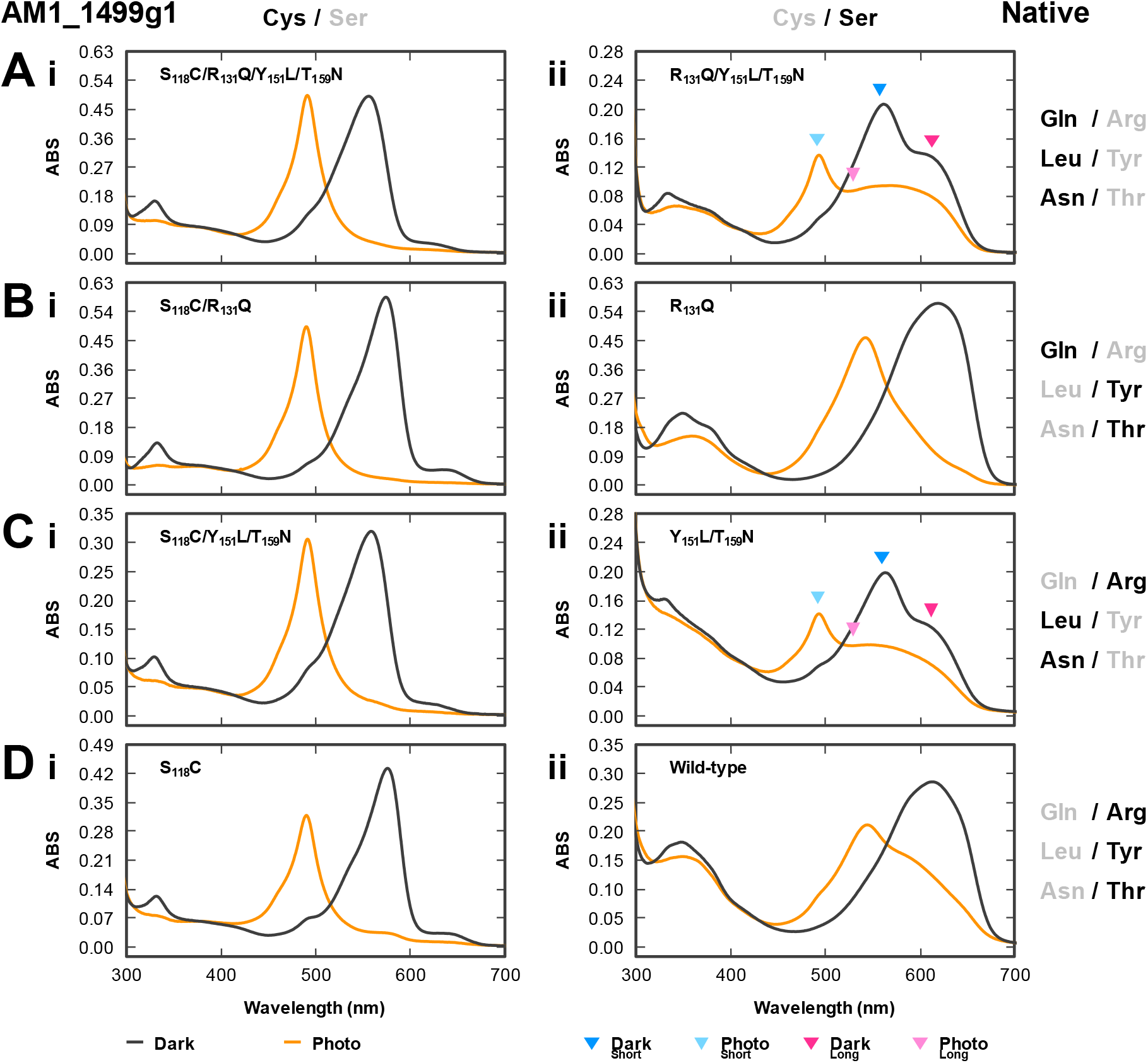
Absorption spectra of native AM1_1499g1 species. (A–D) Absorption spectra of the dark state (solid gray line) and the photoproduct state (solid orange line) were measured at room temperature, except for the wild-type and R_131_Q variant molecules measured at low temperature. In the case of molecules containing two photoconvertible components, both components were fixed to the dark state or the photoproduct state. Wavelength area of the positive (dark state, deep color triangle) and negative (photoproduct state, light color triangle) peaks of the normalized difference spectra from the two photoconvertible components (green/teal one in short wavelength region, cyan; orange/green one in the long-wavelength region, magenta) were assigned by colored triangles. The normalized difference spectra are shown in Figure 6. (A) The molecules having Gln, Leu, and Asn in the Gln/Arg, Leu/Tyr, and Asn/Thr positions, respectively. (B) The molecules having Gln, Tyr, and Thr in these positions. (C) The molecules having Arg, Leu, and Asn in these positions. (D) The molecules having Arg, Tyr, and Thr in these positions. (i, ii) The molecules having Cys (i) or Ser (ii) in the second Cys position.

**Figure 6.**
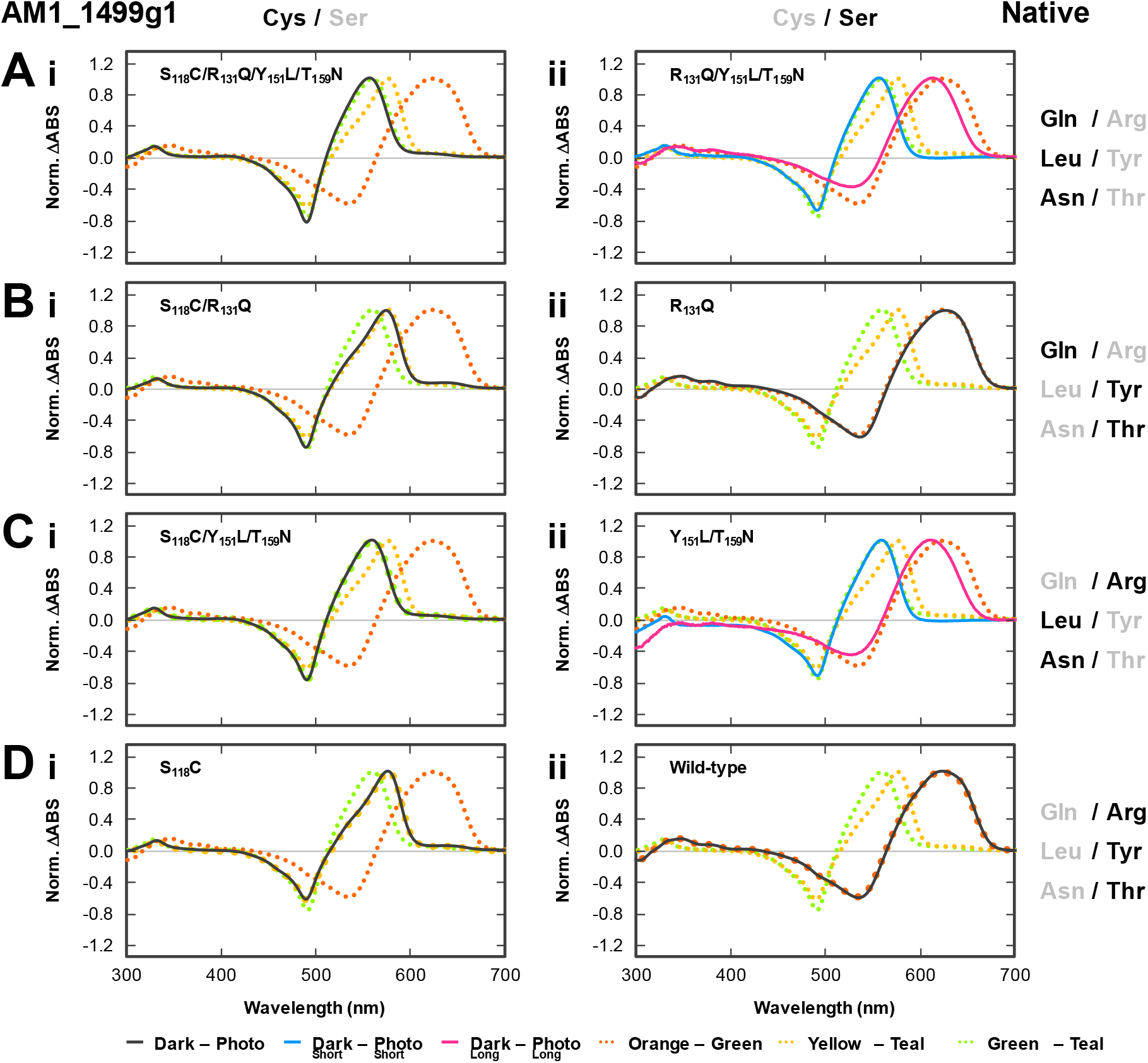
Normalized difference absorption spectra (dark state-photoproduct state) of native AM1_1499g1 species. (A–D) Difference spectra of the molecules containing a single photoconvertible component (solid gray line) were calculated from the native absorption spectra shown in Figure 5. On the other hand, difference spectra of the molecules containing two photoconvertible components (green/teal component in short wavelength region, solid cyan line; orange/green component in the long-wavelength region, solid magenta line) were calculated from the native absorption spectra shown in Figure. S5. In Figure S5, we separately excited each photoconvertible component. These spectra were compared with those of known orange/green, yellow/teal, and green/teal photoconvertible molecules (AM1_1499g1 wild-type, orange dotted line; S_118_C, yellow dotted line; S_118_C/Y_151_L/T_159_N, green dotted line, respectively). These absorption peaks are summarized in Table 1. (A) The molecules having Gln, Leu, and Asn in the Gln/Arg, Leu/Tyr, and Asn/Thr positions, respectively. (B) The molecules having Gln, Tyr, and Thr in these positions. (C) The molecules having Arg, Leu, and Asn in these positions. (D) The molecules having Arg, Tyr, and Thr in these positions. (i, ii) The molecules having Cys (i) or Ser (ii) in the second Cys position.

**Figure 7.**
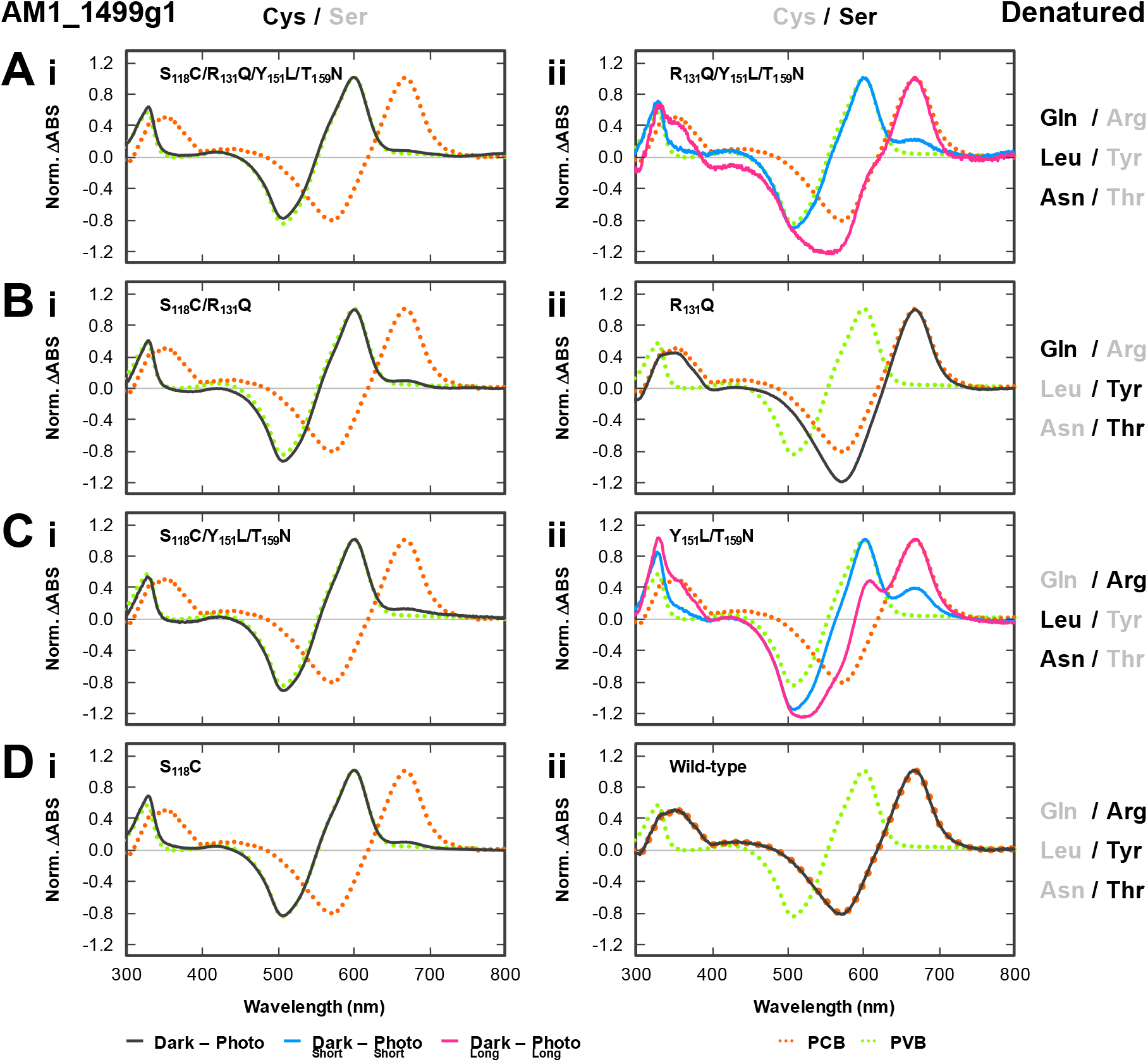
Normalized difference absorption spectra (dark state-photoproduct state) of denatured AM1_6305g1 species. (A–D) Difference spectra of the molecules containing single photoconvertible component (solid gray line) and two photoconvertible components (green/teal component in the short wavelength region, solid cyan line; orange/green component in the long-wavelength region, solid magenta line) were calculated from the denatured absorption spectra shown in Figure S6. Detailed experimental protocol was described in the Materials and Methods. These spectra were compared with those of known PCB- and PVB-binding molecules (AM1_1499g1 wild-type, orange dotted line; AM1_6305g1 wild-type, green dotted line, respectively). These absorption peaks were summarized in Table 1. (A) The molecules having Gln, Leu, and Asn in the Gln/Arg, Leu/Tyr, and Asn/Thr positions, respectively. (B) The molecules having Gln, Tyr, and Thr in these positions. (C) The molecules having Arg, Leu, and Asn in these positions. (D) The molecules having Arg, Tyr, and Thr in these positions. (i, ii) The molecules having Cys (i) or Ser (ii) in the second Cys position.

Under the absence of the second Cys, the phenotypes of the variant molecules were partially inconsistent with the case of AM1_6305g1, particularly in two respects (Fig. 5A-ii to D-ii, Fig. 6A-ii to D-ii, Fig. 7A-ii to D-ii, and Table 1). First, Leu_151_, Asn_159_, and Gln_131_ without the second Cys (R_131_Q/Y_151_L/T_159_N) displayed only partial isomerization activity, harboring both the PCB-binding orange/green component and the PVB-binding green/teal component (Fig. 5A-ii, Fig. 6A-ii, 7A-ii, and Table 1). Second, Gln_131_ had almost no contribution to the isomerization activity because the R_131_Q variant did not show detectable isomerization activity compared to the wild-type (Fig. 5B-ii and D-ii, Fig. 6B-ii and D-ii, 7B-ii and D-ii, and Table 1). Conversely, the functions of the Tyr_151_ and Thr_159_ positions were consistent with the case for AM1_6305g1. Namely, Y_151_L/T_159_N replacement based on the wild-type and R_131_Q backgrounds resulted in the restoration of partial, but not complete, isomerization activity (Fig. 5A-ii and C-ii, Fig. 6A-ii and C-ii, 7A-ii and C-ii, and Table 1). Characterization of the singly mutated variants (Y_151_L and T_159_N) based on the wild-type revealed that the Y_151_L replacement had a greater effect on the isomerization activity than the T_159_N substitute (Fig. S3C, D). The Y_151_L replacement resulted in a significant promotion of isomerization, whereas the T_159_N replacement had a subtle effect on isomerization activity. This differential contribution is consistent with the case of the AM1_6305g1 scaffold (Fig. S3A, B). The dark-state color-tuning is also applicable to the PCB-binding molecules, based on comparison of the difference spectra of the PCB-binding components between the wild-type and Y_151_L/T_159_N molecules; the dark state of the wild-type possessing Tyr_151_ and Thr_159_ was red-shifted in comparison with that of the Y_151_L/T_159_N variant molecule possessing Leu_151_ and Asn_159_ (Fig. 6A-ii, 6C-ii and D-ii, and Table 1). The R_131_Q variant showed a temperature-dependent spectral shift without chromophore isomerization, similar to the wild-type; photoconversion between an orange-absorbing form peaking at 619 nm and a green-absorbing form peaking at 542 nm was observed at a low temperature (5°C), whereas photoconversion between the orange-absorbing form peaking at 628 nm and the green-absorbing form peaking at 545 nm was observed at a high temperature (30°C) (Fig. S4)(Fushimi et al. 2020).

### Possible mechanism of the PCB-to-PVB isomerization

To begin, we discuss the possible PCB-to-PVB isomerization mechanism of the typical DXCF CBCR GAF domains showing blue/green reversible photoconversion, such as TePixJg. A previous reconstitution study has revealed the reaction flow of chromophore incorporation and isomerization (Ishizuka et al. 2011). PCB is initially incorporated into the protein pocket, and the first Cys covalently binds to C3^1^ position of the chromophore by self-ligation activity (Fig. 8A-i). The second Cys ligates to the C10 position of the chromophore in the dark state prior to isomerization from PCB to PVB after chromophore incorporation (Fig. 8A-ii). At this point, blue/green reversible photoconversion has already been observed, albeit with PCB incorporation, suggesting that the C4=C5 double bond of PCB between the A- and B-rings is highly distorted. Furthermore, the C10 bridge between the B- and C-rings would be very bent by the second Cys ligation based on structural information of the PVB-binding TePixJg. As a result, the high distortion of the C4=C5 double bond was stably established by dual fixation of the chromophore at the A-ring and C10 position via first and second Cys residues. Such a highly unstable and rigid conformation would promote the reduction of the C4=C5 double bond and concomitant oxidation of the C2–C3 single bond of the A-ring by a reaction model proposed by Rockwell et al. (Rockwell et al. 2012). After a series of reactions, PCB is finally isomerized to PVB.

**Figure 8.**
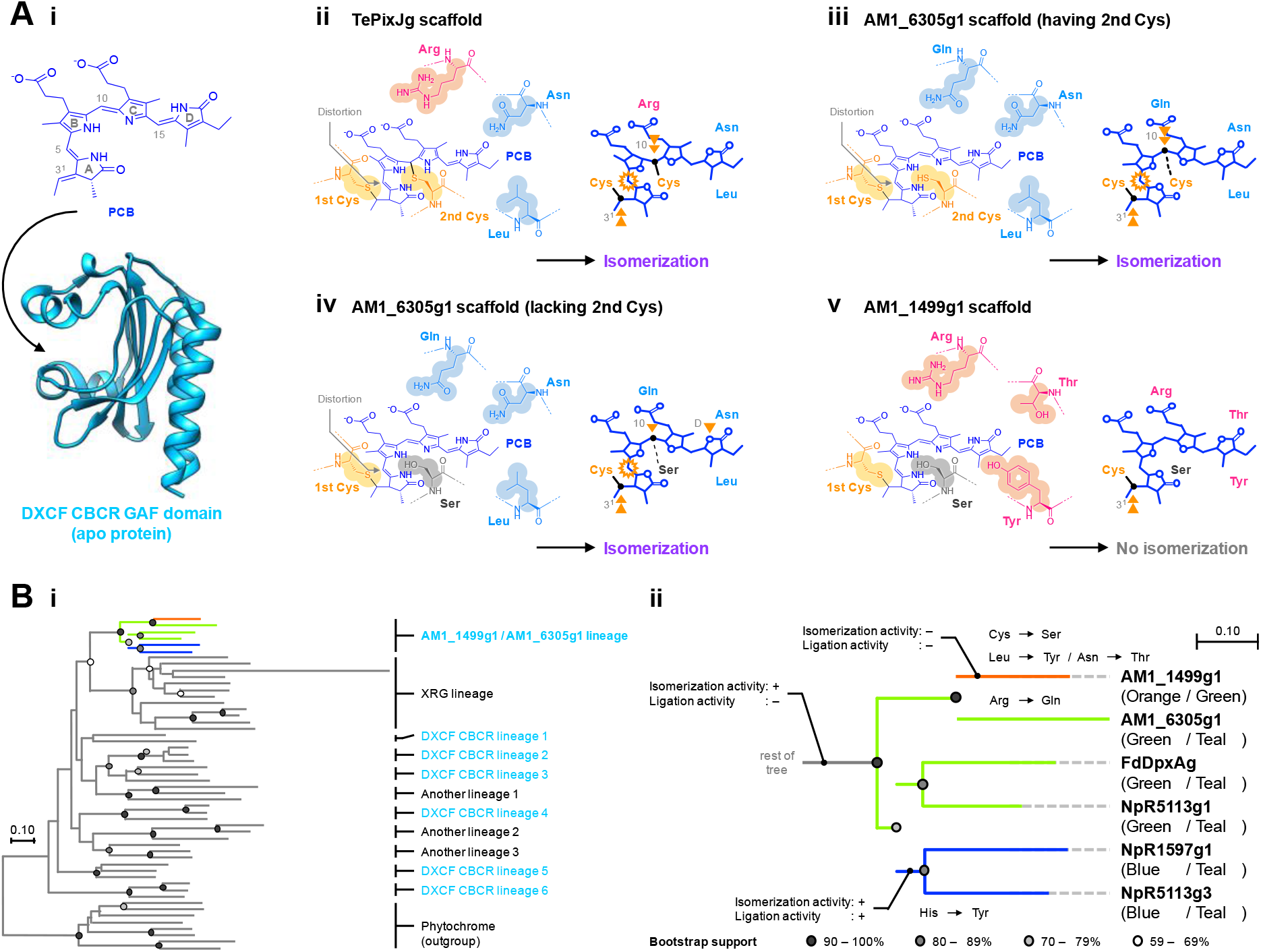
Molecular evolution of DXCF CBCR GAF domains in AM1_1499g1/AM1_6305g1 lineage. (A) Possible molecular mechanisms for PCB-to-PVB isomerization. (i) PCB incorporation into apo DXCF CBCR GAF domain by self-ligation activity. The incorporated chromophore forms a covalent bond between its C3^1^ position and the first Cys of the apo protein. (ii) Interaction network after PCB incorporation in the TePixJg scaffold (typical blue/green photoconvertible molecule which has Arg, Leu, and Asn in the Gln/Arg, Leu/Tyr, and Asn/Thr positions, with the second Cys). The bent PCB is dually fixed at the A-ring and C10 position by the first and second Cys residues with the ligation. This dual fixation mode results in distortion of the C5 bridge between A- and B-rings and promotes PCB-to-PVB isomerization. (iii) Interaction network after PCB incorporation in the AM1_6305g1 scaffold having the second Cys (green/teal photoconvertible molecule which has Gln, Leu, and Asn in these positions, with the second Cys). The unbent PCB is dually fixed at the A-ring and C10 bridge by the first and second Cys residues, although the second Cys does not form covalent bonds. (iv) Interaction network after PCB incorporation in the AM1_6305g1 scaffold lacking the second Cys (C_99_S variant molecule which has Gln, Leu, and Asn in these positions, without the second Cys). The unbent PCB is fixed at the A-ring by the first Cys residue. At the other side, the C10 bridge of the chromophore is weakly held by the Ser residue instead of the second Cys residue, which is supported by the three key residues near the ring D. (v) Interaction network after PCB incorporation in the AM1_1499g1 scaffold (orange/green photoconvertible molecule which has Arg, Tyr, and Thr in these positions, without the second Cys). Because of the lack of neither the second Cys nor the three key residues, no PCB-to-PVB isomerization activity is observed without distortion at the C5 bridge. (B) Phylogenetic tree of diversified CBCR GAF domains with phytochrome GAF domains as the outgroup, which was constructed based on the alignment shown in Figure S7. (i) Each lineage cluster in the whole tree is classified roughly according to photocycle (AM1_1499g1/AM1_6305g1, Asp-Xaa-Cys-Phe (DXCF), expanded red/green (XRG), and the other lineages). CBCR subfamilies possessing the DXCF motif were indicated with cyan. (ii) The AM1_1499g1/AM1_6305g1 lineage is enlarged from the whole tree. Orange, green, and blue lines show the possible emergence and trace of the orange/green, green/teal, and blue/teal CBCR GAF domains, respectively.

On the other hand, although the green/teal CBCR GAF domains such as AM1_6305g1 do not form covalent bond formation between the second Cys and the C10 position of the chromophore in both absorbing forms (Fig. 8A-iii)(Hasegawa et al. 2018), these domains retain the isomerization activity. Therefore, these domains can display the isomerization activity without covalently fixing the C10 of the chromophore. In this context, distortion of the C4=C5 double bond would be established in an alternative way. In this study, we determined that three residues of AM1_6305g1 (Gln_112_, Leu_132_, and Asn_140_) contributed to isomerization activity when the second Cys was replaced with Ser (Fig. 1A-ii to D-ii, Fig. 2A-ii to D-ii, Fig. 3A-ii to D-ii, and Table 1). Conversely, these residues are dispensable under the presence of the second Cys (Fig. 1A-i to D-i, Fig. 2A-i to D-i, Fig. 3A-i to D-i, and Table 1). The second Cys, but not the Ser, can completely hold the chromophore in the C4=C5 distorted conformation. The thiol group of the second Cys may have a unique electrostatic interaction with the chromophore near the C10 position, which could not be fully complemented by the hydroxy group of the Ser.

Under the absence of the second Cys, Gln_112_/Leu_132_/Asn_140_ residues in addition to Ser_99_, cooperatively work to produce the isomerization activity (Fig. 8A-iv). Ser_99_ may have a weaker interaction with the chromophore than the second Cys, and these three residues would support the function of Ser_99_. The Leu_132_/Asn_140_ residues have a larger contribution than the Gln_112_ residue (Fig. 1A-ii to D-ii, Fig. 2A-ii to D-ii, Fig. 3A-ii to D-ii, and Table 1). The Leu_132_/Asn_140_ residues have been elucidated to fix the D-ring into a twisted geometry, which would assist the Ser_99_ in the distortion of the C4=C5 double bond. Gln_112_ is predicted to be positioned near the C-ring propionate and contribute to the stabilization of the chromophore conformation.

However, this prediction is based on the TePixJg structure with the “bent” chromophore, whose conformation would be largely different from that of “unbent” one (Fig. S1B, D). Therefore, it is difficult to elucidate the precise role of Gln_112_ in AM1_6305g1. It is of note that not only does the second Cys have no direct interactions with the C4=C5 double bond, but the three residues also do not. In particular, these three residues would have interaction with the chromophore around the C- and D-rings far from the C4=C5 bond. Fixation of the chromophore on both sides of the C5 bridge between the A- and B-rings would be important for isomerization activity, even though one side of the fixation site is far from this bridge with another side fixed at C3^1^ by the first Cys. Although no naturally occurring molecules showing isomerization activity in the absence of the second Cys have been reported, this study provides the potential for the presence of such natural molecules that apply a remote fixation system.

### Evolutionary trace of the AM1_1499g1/AM1_6305g1 lineage

We constructed a phylogenetic tree of the CBCR GAF domains with the phytochrome GAF domains as the outgroup for the evolutionary trace (Fig. 8B-i). The DXCF CBCR GAF domains are detected in several lineages, and the AM1_1499g1/AM1_6305g1 lineage is one of such families. This lineage shares an ancestor with the expanded red/green (XRG) lineage GAF domains. The first branching resulted in two subgroups within this lineage (Fig. 8B-ii). Notably, these two subgroups include green/teal photoconvertible molecules, indicating that this lineage’s ancestral molecule would be the green/teal photoconvertible version showing isomerization activity but not reversible Cys ligation capabilities. Furthermore, molecules of this lineage, except for AM1_1499g1, retain the Leu/Asn residues that largely contribute to the isomerization activity without the second Cys residue (Fig. S1E). We consider that, because the ancestral molecule of this lineage originally lost the ligation ability conferred by the second Cys, the weaker interaction mode of the second Cys has promoted the acquisition of the Leu/Asn residues for robust isomerization activity.

These subgroups are further divided into two populations. Blue/teal photoconvertible molecules showing not only isomerization activity but also reversible Cys ligation have emerged in the case of one subgroup (Fig. 8B-ii). This event would occur with the replacement of the His residue with Tyr next to the first Cys, in which the bulky Tyr side chain may push the C10 bridge to facilitate Cys ligation at the opposite side, based on the previous study (Fig. S1E)(Fushimi et al. 2020). On the other hand, in the case of the other subgroup, AM1_1499g1 has emerged, which has lost the second Cys and does not show isomerization activity (Fig. 8B-ii and Fig. S1E). However, the present study identified that the replacement of only the second Cys is insufficient for the complete inactivation of the isomerization activity. In fact, AM1_1499g1 has lost not only the second Cys but also the Leu/Asn residues (Fig. S1E). AM1_1499g1 has experienced multiple amino acid replacements after diversification to establish a longer wavelength perception (Fig. 8A-v and B). On the other hand, AM1_6305g1 has specifically acquired Gln_112_, which supports isomerization activity for a more robust isomerization system (Fig. 8B-ii and Fig. S1E). The whole protein domain architectures are quite similar to each other and these proteins are derived from the same organism, *Acaryochloris marina*, suggesting that these two domains may have evolved in the opposite direction to acquire dual windows for shorter and longer light perception after the protein duplication event. In summary, molecules within this lineage have undergone unique evolutionary events to develop highly diverse green/teal, blue/teal, and orange/green photocycles through only five amino acid replacements in total.

## Materials and Methods

### Bacterial strains and growth media

The *Escherichia coli* strain JM109 (TaKaRa) was used for plasmid DNA cloning. The *E. coli* strain C41 (Cosmo Bio) harboring plasmid pKT271 (constructed by heme oxygenase and phycocyanobilin:ferredoxin oxidoreductase genes from *Synechocystis* sp. PCC 6803)(Mukougawa et al. 2006) was used for co-expression of proteins with the PCB-producing system. Bacterial cells were grown on lysogeny broth (LB) agar medium containing 20 μg/mL kanamycin with or without 20 μg/mL chloramphenicol for selection of plasmid-transformed cells.

### Bioinformatics

Multiple sequence alignment and neighbor-joining phylogenetic trees were constructed using MEGA7 software (Kumar, Stecher, and Tamura 2016). The crystal structure of PVB–bound TePixJg (blue-absorbing form, PDB_ID: 4GLQ) was utilized to assess key amino acid residues for isomerization. The molecular graphic was generated using the UCSF Chimera software (Pettersen et al. 2004).

### Plasmid construction

AM1_6305g1 (amino acid positions 33–203)(Hasegawa et al. 2018) and AM1_1499g1 (amino acid positions 47–222)(Fushimi et al. 2020) genes fused with an N-terminal His-tag sequence have been inserted into a pET28a vector (Novagen) for protein expression in *E. coli.* The plasmids of AM1_6305g1 and AM1_1499g1 variants were generated by site-directed mutagenesis using each parent plasmid. KOD One PCR Master Mix (Toyobo Life Science) with the appropriate nucleotide primer sets was used for mutagenesis. The primer sets of forward primer GATGATxxxTTTGGTCATAATTATGCCAATAAA and reverse primer ACCAAAyyyATCATCGCGTACATTGGC for the replacement of Cys_99_ of AM1_6305g1 with Ser (xxx = agt, yyy = act) and Ala (xxx = gcc, yyy = ggc) were prepared. The primer set of forward primer CAAGGAcggGTGTTTGCCGTTGCGGAT and reverse primer AAACACccgTCCTTGCTGATATTTATTGGC for the replacement of Gln_112_ of AM1_6305g1 with Arg, and conversely, the primer set of forward primer TTGGGGcagGTATTTGCAGTAGCCGATGTCTACGAT and reverse primer AAATACctgCCCCAATTGATATTTGGTCGCATAATT for the replacement of Arg_131_ of AM1_1499g1 with Gln were prepared, respectively. The other primer sets for the replacements of Leu_132_ of AM1_6305g1 with Tyr, Asn_140_ of AM1_6305g1 with Thr, Ser_118_ of AM1_1499g1 with Cys, Tyr_151_ of AM1_1499g1 with Leu, and Thr_159_ of AM1_1499g1 with Asn were constructed in a previous study (Fushimi et al. 2020). All plasmid sequences were confirmed by DNA sequencing (Eurofins Genomics).

### Protein expression and purification

*E. coli* C41 pKT271 was cultured in 1 L LB medium at 37°C until the optical density at 600 nm was 0.4–0.8, and then the cells were cultured overnight at 18°C after isopropyl β-D-1-thiogalactopyranoside (IPTG) addition (final concentration, 0.1 mM) for induction of the protein expression. The cells were collected by centrifugation at 5,000 g for 15 min and were suspended in lysis buffer (20 mM HEPES-NaOH pH 7.5, 0.1 M NaCl, and 10% (w/v) glycerol). All proteins were extracted and purified as described in the previous study (Fushimi et al. 2020).

### SDS-PAGE analysis

Purified proteins were diluted in buffer (60 mM Tris–HCl pH 8.0, 2% (w/v) sodium dodecyl sulfate (SDS), and 60 mM dithiothreitol (DTT)), denatured at 95°C for 3 min, and electrophoresed at room temperature (r.t.) using 12% (w/v) SDS polyacrylamide gels. The electrophoresed gels were soaked in distilled water for 30 min and then exposed to blue light (λ_max_ 470 nm) and green light (λ_max_ 527 nm) using a WSE-5500 VariRays (ATTO) with a short-path filter (passing through < 562 nm) to visualize the fluorescence of the proteins. The fluorescence bands were imaged using a WSE109 6100 LuminoGraph (ATTO) with a long-path filter (passing through > 600 nm). After observation, the gels were stained with Coomassie brilliant blue R-250 (CBB).

### UV-Vis spectroscopic analysis

A UV-2600 spectrophotometer (SHIMADZU) was used to monitor the ultraviolet and visible absorption spectra of the proteins. An Opto-Spectrum Generator (Hamamatsu Photonics) was used to prepare monochromatic light of various wavelengths to induce photoconversion: teal-absorbing form, 470–490 nm; green-absorbing form, 490–580 nm; yellow-absorbing form, 580–620 nm; orange-absorbing form, 620–640 nm. Among the AM1_6305g1 and the AM1_1499g1 variants, the molecules containing both green or yellow/teal and orange/green photoconvertible components were irradiated with each monochromatic light independently for conversion into the dark state (15*Z*–isomer) and the photoproduct state (15*E*–isomer). The absorption spectra of these native proteins were recorded at r.t., except for the AM1_1499g1 wild-type and R_131_Q variants, which were measured at low (5°C) and high (30°C) temperatures. Normalized difference absorption spectra (“dark state–photoproduct state” of either photoconvertible components) were calculated from these raw absorption spectra (shown in Fig. S5). Furthermore, the normalized double-difference spectrum (normalized difference spectrum of the yellow/teal component–normalized difference spectrum of the orange/green component) of AM1_1499g1_T_159_N was calculated from these normalized difference absorption spectra. Photocycle forms were decided by comparing these spectra between each variant molecule and known orange/green, yellow/teal, and green/teal photoconvertible molecules (AM1_1499g1 wild-type, S_118_C and S_118_C/Y_151_L/T_159_N, respectively)(Fushimi et al. 2020).

### Identification of the chromophore incorporated into the AM1_6305g1 and AM1_1499g1 variant molecules containing two photoconvertible components

We established the experimental protocol in this study to determine the binding chromophore species of the molecules containing both green-to-yellow/teal and orange/green photoconvertible components. We first prepared the sample in which one photoconvertible component was fixed in the photoproduct state (15*E*-isomer), while another photoconvertible component was fixed to the dark state (15*Z*-isomer). The sample under such a condition was subjected to denaturation with 8 M acidic urea (pH < 2.0). Irradiation of the sample with white light (3 min) caused photoconversion of the 15*E*-isomer of one component. We determined the binding chromophore species of each photoconvertible component based on comparison of the difference spectrum after and before white light illumination (shown in Fig. S6) with those of the PCB-binding and PVB-binding reference molecules (AM1_1499g1 and AM1_6305g1, respectively)(Hasegawa et al. 2018; Fushimi et al. 2020). Furthermore, the normalized double-difference spectra (normalized difference spectrum of the yellow/teal component–normalized difference spectrum of the orange/green component) of AM1_6305g1_C_99_S/L_132_Y/N_140_T and C_99_S/Q_112_R/L_132_Y/N_140_T were calculated from these normalized difference absorption spectra.

## Acknowledgments

We thank Prof. J. Clark Lagarias and Dr. Nathan C. Rockwell (University of California, Davis) for helpful discussion, and Enago (https://www.enago.jp/) for the English language review. This work was supported by Japan Science and Technology Agency, Core Research for Evolutional Science and Technology (JPMJCR1653, to R.N.).

---

**Figure S1.**
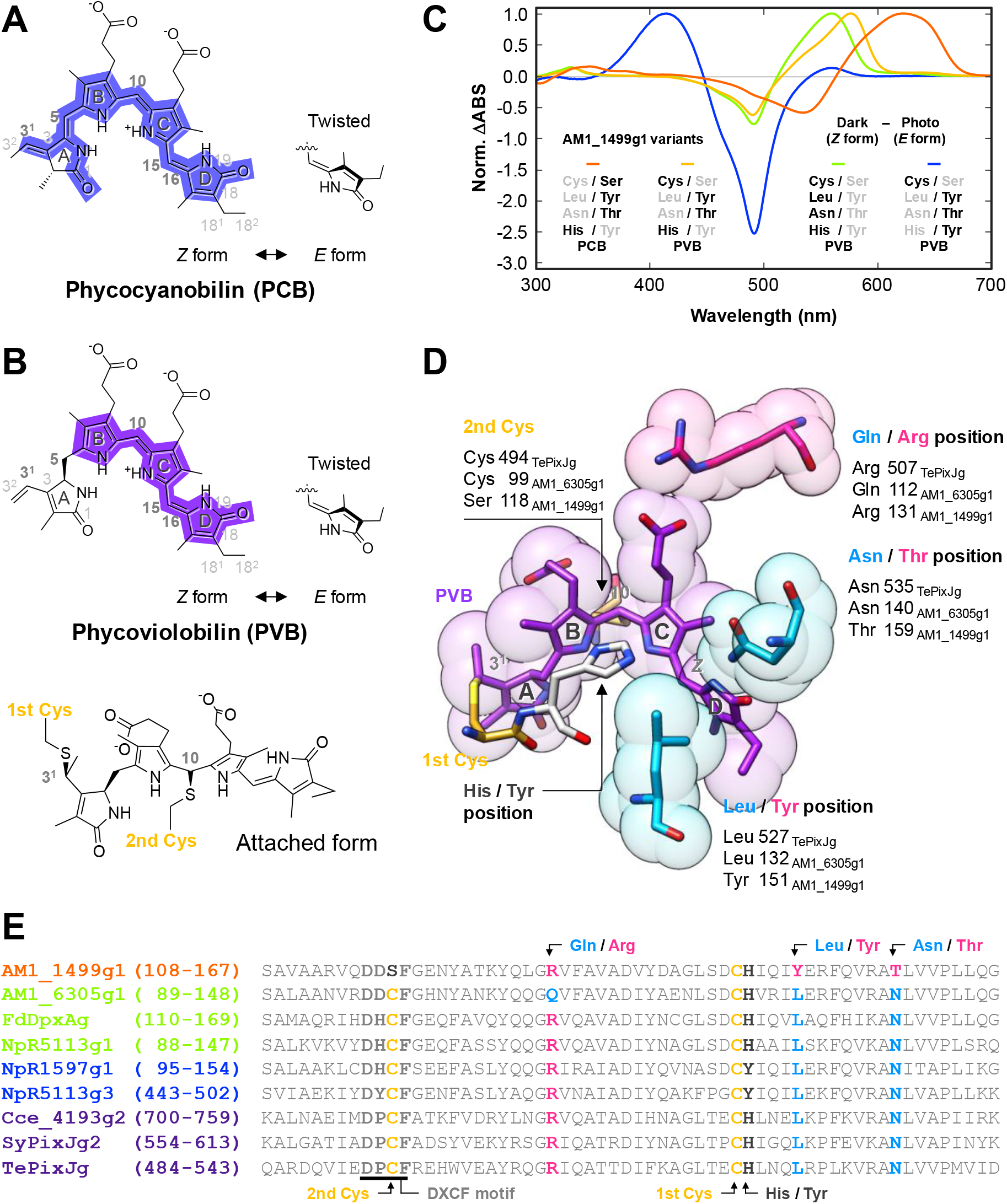
Diversity of DXCF CBCR GAF domains. (A, B) Chemical structures of phycocyanobilin (PCB) (A) and phycoviolobilin (PVB) (B). These structures are shown in their *15Z* dark states, with π–conjugated systems highlighted. Photoisomerization of the C15=C16 double bond yields the 15*E* photoproduct states, for which the D-rings are shown in twisted conformation. Furthermore, the second Cys attached form of PVB in the *15Z* dark state was shown. (C) Normalized difference spectra (*15Z* form I5*Λ*’ form) of AM1_1499g1 variant molecules (orange, wild-type showing orange/green photocycle; yellow, S_118_C variant showing yellow/teal photocycle; green, S_118_C/Y_151_L/T_159_N variant showing green/teal photocycle; blue, S_118_C/H_147_Y showing blue/teal photocycle). (D) Key residues involved in the isomerization and reversible ligation are shown on the crystal structure of DXCF CBCR GAF domain (TePixJg in the blue-absorbing form with *15Z–PVB* (purple)(PDB ID 4GLQ))(Burgie et al. 2013). The coloring of these residues corresponds to that of residues in the multiple alignment in Figure S1E. (E) Multiple sequence alignment of diversified CBCR GAF domains, which show orange/green (orange), green/teal (green), blue/teal (blue), and blue/green (purple) photocycles. Specific residues on these sequences were highlighted; the first Cys and the second Cys in the DXCF motif (yellow), His/Tyr position (gray), and three residues in the Leu/Tyr, Asn/Thr, and Gln/Arg positions characterized in AM1_6305g1 (cyan) and AM1_1499g1 (magenta).

**Figure S2.**
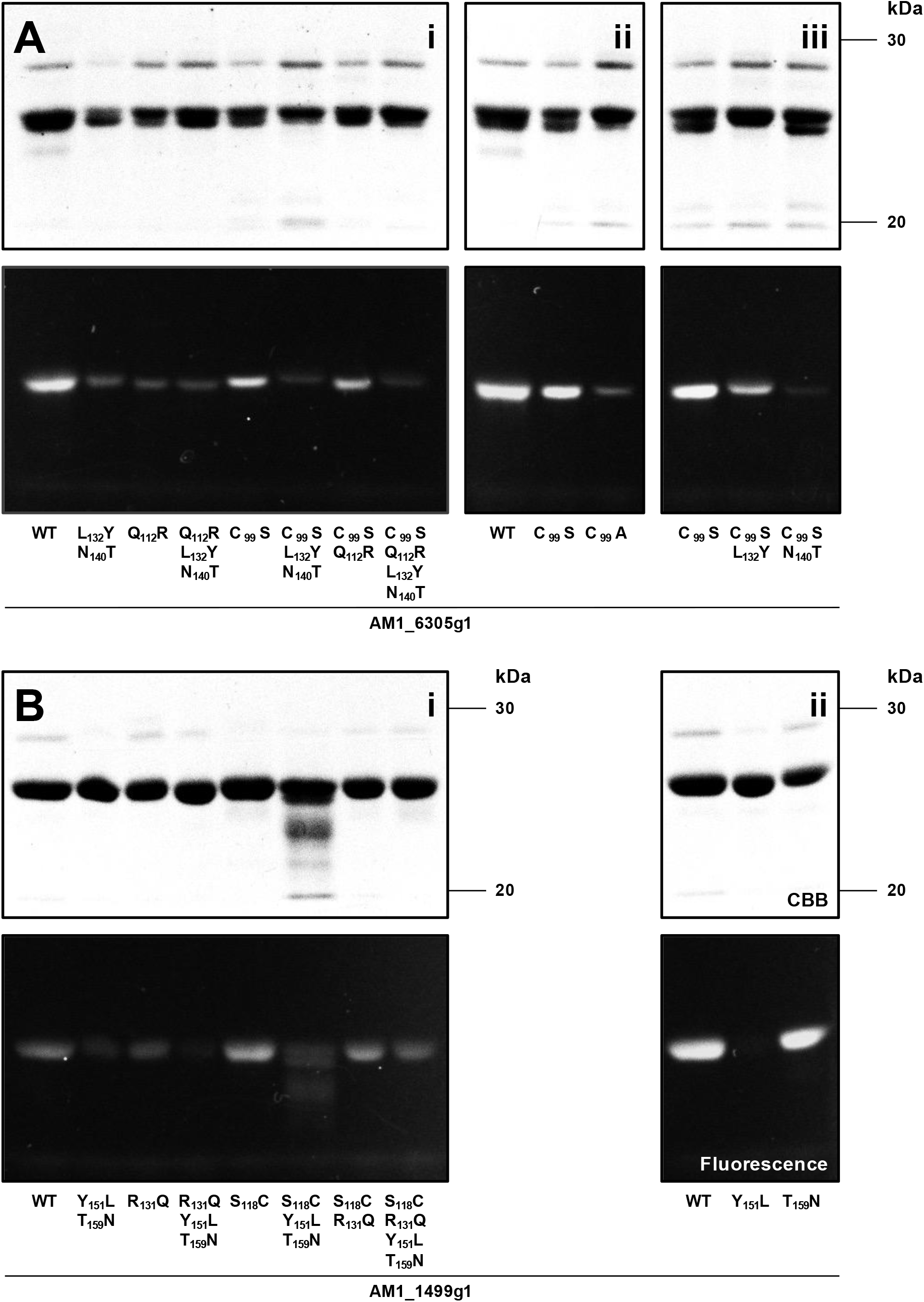
Purified AM1_6305g1 and AM1_1499g1 variant molecules. (A, B) SDS-PAGE analyses of the AM1_6305g1 (A) and AM1_1499g1 (B) species. Coomassie Brilliant Blue (CBB) stained gels (upper) and fluorescence imaging gels (lower).

**Figure S3.**
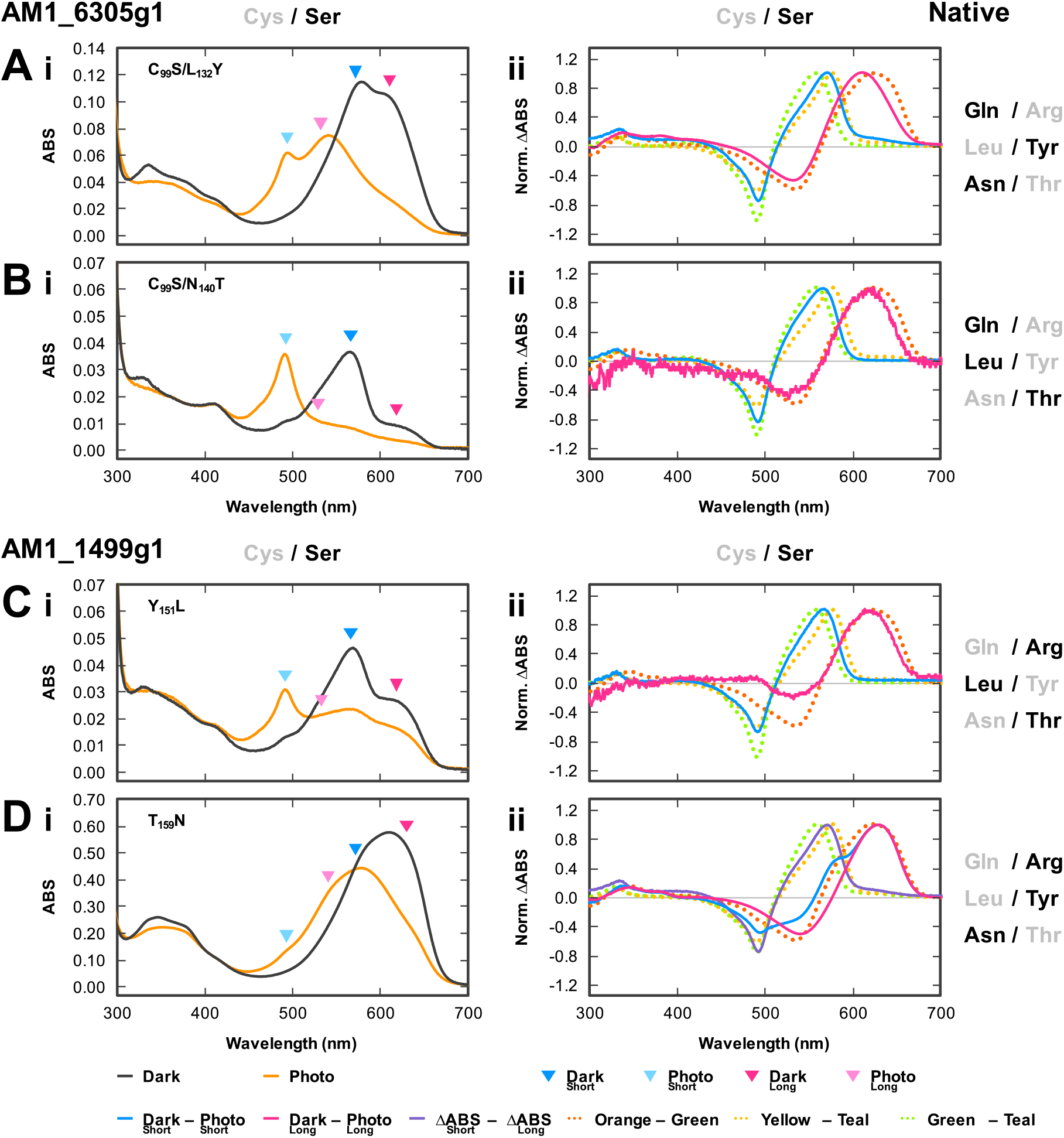
UV-Vis spectra of native AM1_6305g1 and AM1_1499g1 species. (A-i to D-i, left column) The absorption spectra (dark state, solid gray line; photoproduct state, solid yellow line) of AM1_6305g1 (A-i, B-i) and AM1_1499g1 (C-i, D-i) variant molecules were measured at room temperature, in which both components were fixed to the dark state or the photoproduct state. Wavelength area of the positive (dark state, deep color triangle) and negative (photoproduct state, light color triangle) peaks of the normalized difference spectra from the two photoconvertible components (green-to-yellow/teal component in the short wavelength region, cyan; orange/green component in the long-wavelength region, magenta) were assigned by colored triangles. (A-ii to D-ii, right column) The normalized difference spectra (dark state–photoproduct state) of AM1_6305g1 (A-ii, B-ii) and AM1_1499g1 (C-ii, D-ii) variant molecules containing the two photoconvertible components (green-to-yellow/teal component in short wavelength region, solid cyan line; orange/green component in the long-wavelength region, solid magenta line) were calculated from the native absorption spectra shown in Figure S5. Furthermore, the normalized double-difference spectrum (yellow/teal photoconvertible component in the short wavelength region, solid violet line) of AM1_1499g1_T_159_N was calculated from these normalized difference absorption spectra. These spectra were compared with those of the known orange/green, yellow/teal, and green/teal photoconvertible molecules (AM1_1499g1 wild-type, orange dotted line; S_118_C, yellow dotted line; S_118_C/Y_151_L/T_159_N, green dotted line, respectively).

**Figure S4.**
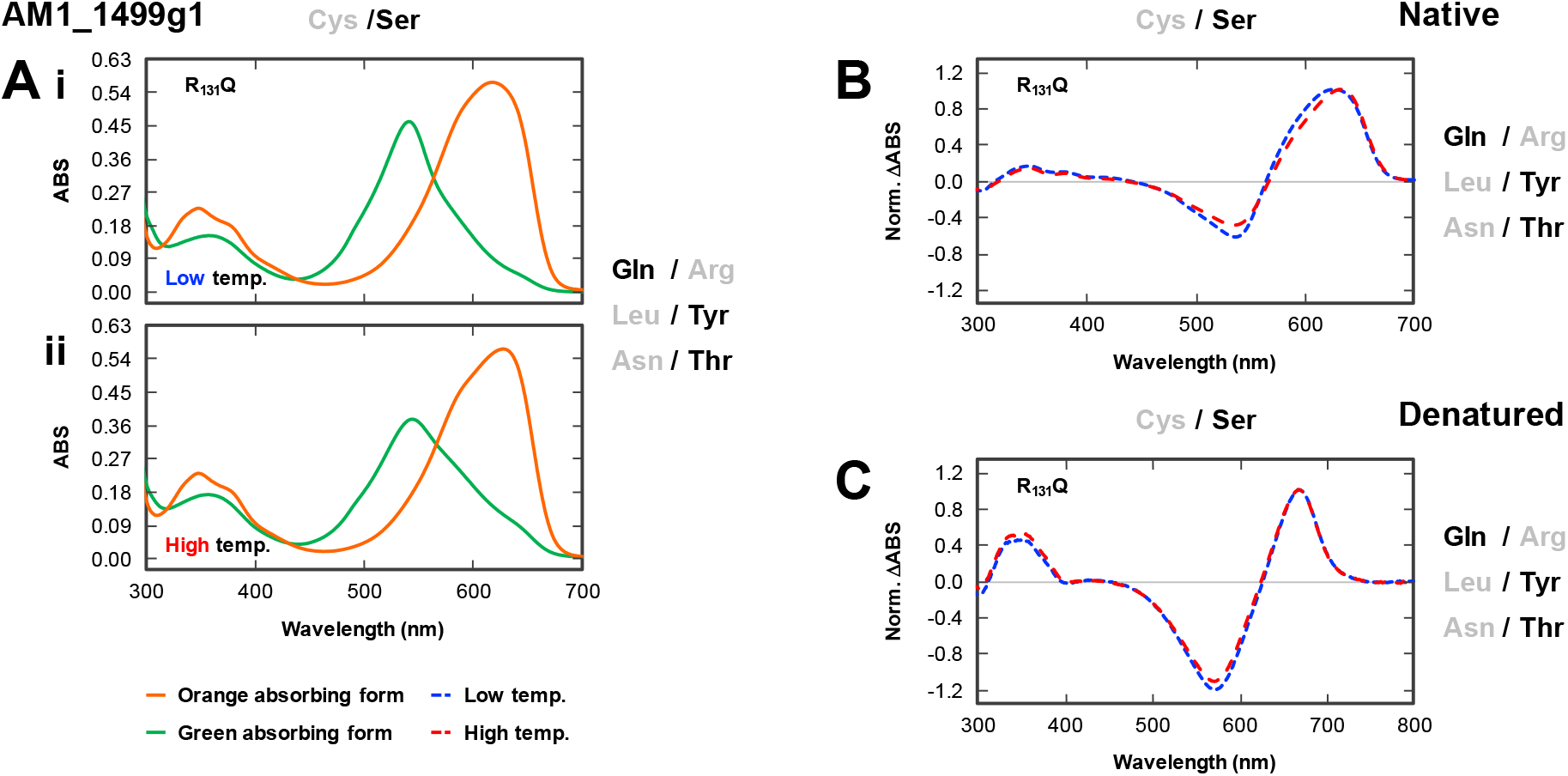
Temperature-dependent photocycle of AM1_1499g1_R_131_Q. (A) Absorption spectra of AM1_1499g1_R_131_Q variant at low (5°C) (i) and high (30°C) (ii) temperature (orange-absorbing form, solid orange line; green-absorbing form, solid green line). (B, C) Normalized difference spectra (dark state–photoproduct state) of the native (B) and denatured (C) proteins at 5°C (blue dashed line) and 30°C (red dashed line).

**Figure S5.**
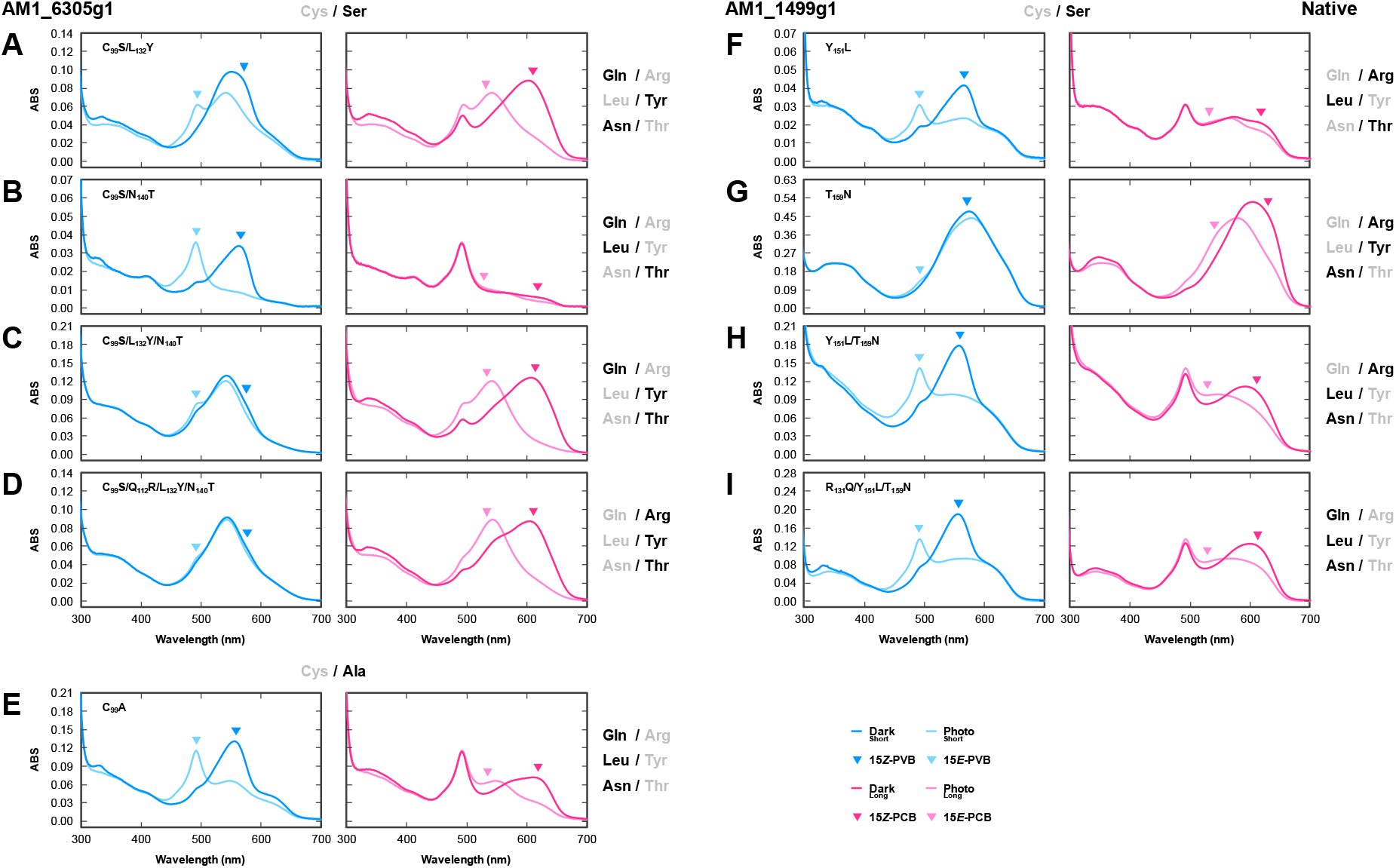
Photoconversion of one of the two photoconvertible components of the AM1_6305g1 and AM1_1499g1 variant molecules. The green/teal or yellow/teal (cyan) and orange/green (magenta) photoconvertible components contained in the AM1_6305g1 (A–E) and AM1_1499g1 (F–I) variant molecules were independently excited by specific monochromic light sources to observe each photoconversion between the dark state (15*Z*–isomer, deep color) and the photoproduct state (15*E*–isomer, light color). The difference spectra shown in Figures 2, 4, 6, and S3 were calculated from these absorbance spectra.

**Figure S6.**
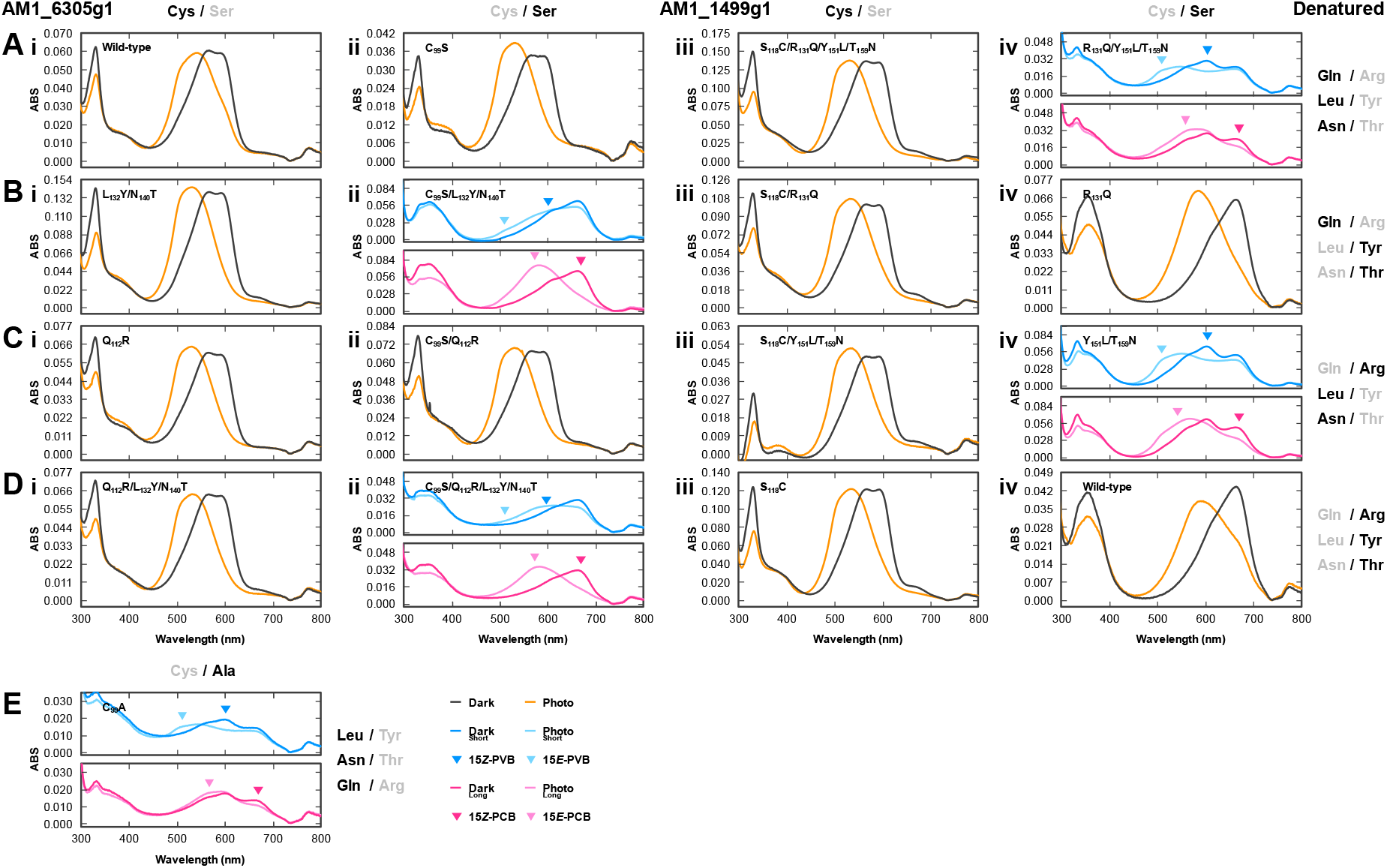
Absorption spectra of denatured AM1_6305g1 and AM1_1499g1 species. (A–E) In the case of the molecules containing a single photoconvertible component, the photoproduct state (15*E*–isomer, yellow) was denatured by acid urea and was converted into the dark state (15*Z*–isomer, gray) by white light illumination. On the other hand, in the case of molecules containing two photoconvertible components, we first prepared the sample in which one photoconvertible component ((green or yellow)/teal (cyan) or orange/green (magenta) component) was fixed to the photoproduct state (15*E*-isomer, light color), while another photoconvertible component was fixed to the dark state (15*Z*-isomer, deep color), which was subjected to the acid denaturation analysis. Details are described in the Materials and Methods. The difference spectra shown in Figures 3, 4, and 7 were calculated from these denatured spectra. (F) Model of the acid denaturation assay. The chromophore incorporated into native proteins can reversibly photoconvert by appropriate light irradiation before acid-urea treatment. Conversely, the chromophores covalently bound to denatured proteins irreversibly isomerize to 15*Z*–form by light irradiation after acid-urea treatment.

**Figure S7.**
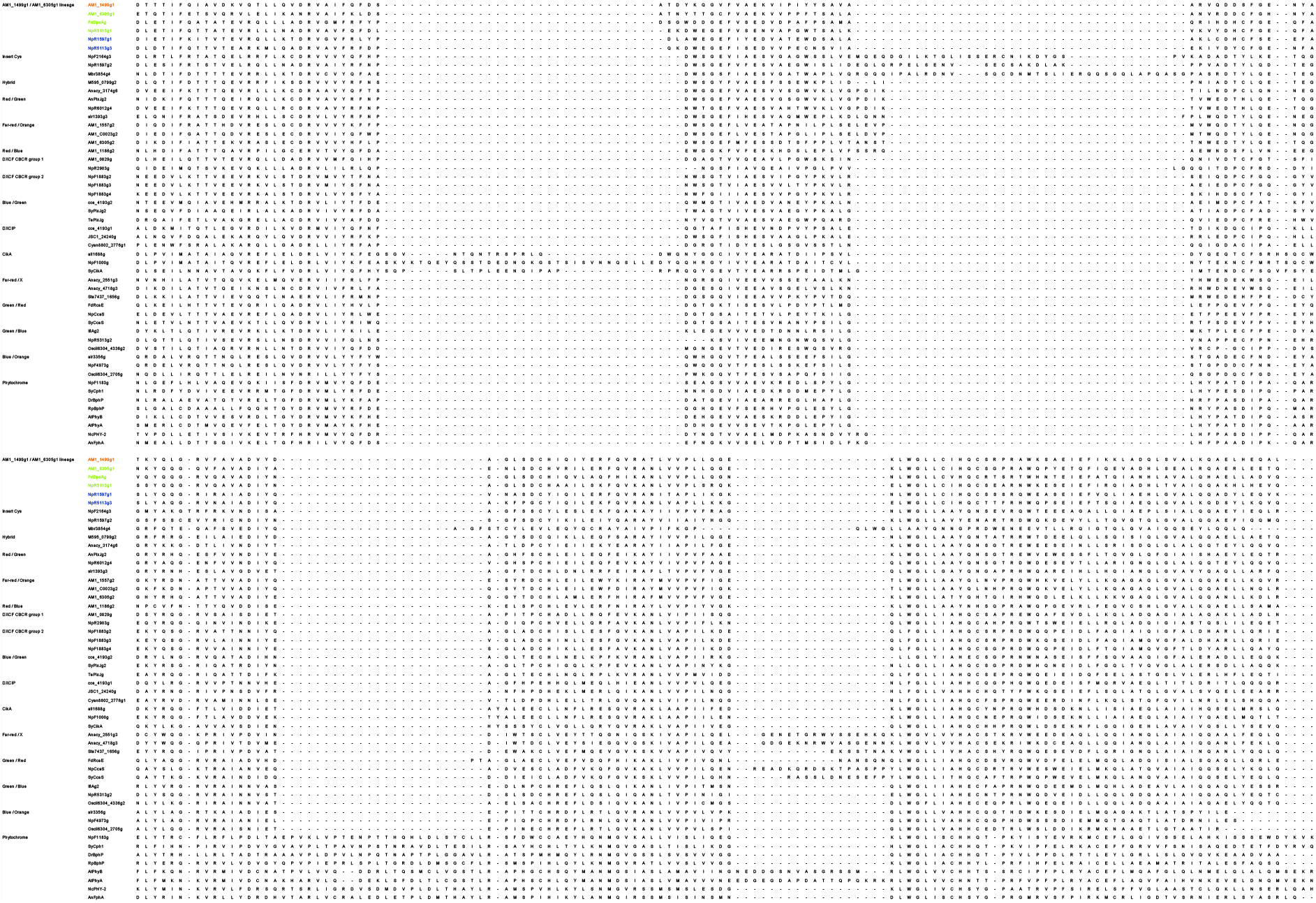
Multiple sequence alignment of diversified CBCR GAF domains. CBCR GAF domains in AM1_1499g1/AM1_6305g1 lineage were categorized as orange/green (orange), green/teal (green), and blue/teal (blue) photoconvertible molecules. The phylogenetic tree shown in Figure 8B was constructed from this alignment.

